# Tracking microbial evolution in the human gut using Hi-C

**DOI:** 10.1101/594903

**Authors:** Eitan Yaffe, David A. Relman

## Abstract

Despite the importance of horizontal gene transfer for rapid bacterial evolution, reliable assignment of mobile genetic elements to their microbial hosts in natural communities such as the human gut microbiota remains elusive. We used Hi-C (High-throughput chromosomal conformation capture), coupled with probabilistic modeling of experimental noise, to resolve 88 strain-level genomes of distal gut bacteria from two subjects, including 12,251 accessory elements. Comparisons of 2 samples collected 10 years apart for each of the subjects revealed extensive *in situ* exchange of accessory elements, as well as evidence of adaptive evolution in core genomes. Accessory elements were predominantly promiscuous and prevalent in the distal gut metagenomes of 218 adult subjects. This work provides a foundation and approach for studying microbial evolution in natural environments.

---

One of the major forces shaping the genomic landscape of microbial communities is horizontal gene transfer (HGT)^1^. HGT is of particular importance for the human gut microbiome, where it is involved in the emergence of antibiotic-resistant bacterial strains and mobilization of virulence factors^2, 3^. In comparison to other microbial communities, human and other animal gut microbiotas show evidence of especially widespread HGT among bacterial members^4^. Moreover, there is mounting evidence of HGT between bacterial pathogens and commensals, based on *in vitro* experiments^5^ and animal models^6–8^. Because strains can persist for decades within the same subject^9^, the human gut microbiota has the potential to reveal quantitative and time-resolved aspects of HGT in a natural setting, with implications for both microbial evolution and human health.

The genome of any specific microbe is a mosaic of components that follow distinct evolutionary paths, ranging from tightly coupled, co-evolving house-keeping genes, to a collection of loosely associated mobile elements, including bacteriophages, transposons, plasmids, and other non-essential genes^10^. Comparisons of closely related genomes for most generalist microbial species (representing strains of the same species) identify a set of genes that are shared by all strains (‘core’), and a remaining set that are present in only a subset of strains (‘accessory’). These accessory genes contribute to the genetic diversity of the species and the capacity for adaptation to new environmental challenges and conditions^11^. Computational methods based on gene co-occurrence patterns across individuals have identified core genomes from human gut metagenomic data; however, linkage of accessory elements with their hosts has been limited to simple cases of species-specific elements, such as narrow-host-range bacteriophages^12^.

*De novo* genotyping of microbial communities with a complex population structure, such as the human gut microbiota, is challenging for several reasons. First, a community may contain multiple conspecific strains^13^. Second, promiscuous mobile elements may be harbored by multiple microbial hosts in the same community^14, 15^. These features of the genomic landscape prevent robust recovery of genomes from complex communities using standard approaches, such as metagenomic binning^16^. Thus, while core genomes can be inferred from metagenomic data with current methods, characterization of mobile elements and their linkage to host species in natural settings remains elusive.

Hi-C is a fixation-based method for estimating the probability of close physical proximity between DNA fragments^17, 18^. A single Hi-C assay typically produces millions of ‘contacts’, where each contact reflects two sequence fragments that were adjacent in three-dimensional space at the time of fixation. Hi-C maps have revealed large-scale chromatin structures involved in genome regulation in eukaryotes^19, 20^. More broadly, the technique has been used to study DNA folding across the tree of life, from bacteria to mammals^21–23^, and to perform *de novo* genome assembly of isolated species^24–27^. When applied to microbial communities (‘metagenomic Hi-C’), the global nature of Hi-C enables the study of multiple genomes simultaneously. Hi-C has enhanced genome co-assembly, as shown with synthetic bacterial communities^28^, and has facilitated the association of extra-chromosomal DNA with the chromosomes of their microbial hosts^29^. Hi-C has provided insights into virus-host interactions in the mouse gut^30^ and resolved diverse microbial genomes in the human gut^31, 32^. However, both the presence of noise, in the form of spurious inter-cellular contacts, and the potential within-host sharing of genetic elements, have not been adequately addressed thus far with metagenomic Hi-C, confounding the interpretation of the data.

Here we couple metagenomic Hi-C with rigorous probabilistic noise modeling, to genotype the human gut microbiome. Application of the method to samples from two individuals recovered 88 genomes, with accessory genes on average accounting for a quarter of each genome. Analysis of samples collected ten years apart from each of the subjects identified a total of 12 genomes with evidence of within-host strain evolution. A comprehensive analysis of both gene-content and nucleotide-level changes in these 12 strains revealed highly dynamic accessory genomes, along with evidence for adaptive evolution in core genomes. Finally, the majority of the accessory elements identified in the two subjects were prevalent in gut metagenomes of 218 additional adult subjects, where they showed promiscuous associations with multiple strains and species.

## RESULTS

Stool was collected from a healthy adult (subject A); DNA was extracted, paired-end sequenced, and the resulting 202M (million) paired reads were compiled into a metagenome assembly (N50 measure of 4.7Kb), composed of 308K (thousand) contigs (consensus DNA regions) that collectively spanned 648Mb. The same sample was assayed in triplicate using the Hi-C protocol as described in Marbouty et al.^31^, with minor adaptations (**Materials and Methods**). Briefly, stool was treated with formaldehyde, and cells were lysed. DNA was digested using the restriction enzyme DpnII, ligated under dilute conditions using T4 ligase, sheared and size-selected (>500bp), and paired-end sequenced with 1.4B (billion) Hi-C read pairs in total. After quality filtering, 797M read pairs were mapped successfully back onto the assembly. Within contigs, the density of mapped reads varied inversely with the genomic distance between the two paired ends, confirming that the global and stochastic nature of Hi-C data was recapitulated in our system (**fig. S1**). Technical replicates were correlated (Spearman coefficient between inter-contig read count matrices was >0.72) and were therefore united. Downstream analysis was limited to 37.5M inter-contig read pairs (5.6% of total reads). By locating nearby DpnII restriction sites, each read pair was converted into a *contact*, which is a pair of restriction fragment ends that were inferred to have been ligated during the procedure. The resulting contact map contained 10.3M unique inter-contig contacts.

### Genotyping microbial communities using Hi-C

To tackle the complexity of natural microbial communities, we first considered the possible relationships between assembled contigs and microbial strains. We use the term *genome configuration* to refer to a set of contigs that represent the genomic capacity (including extra-chromosomal DNA) of a clonal strain (**Supplementary Text**). In a community composed of distantly related strains that do not exchange genes, there is a one-to-one mapping between strains, configurations and genomes, as contigs are unambiguously related to a single population and genome. The relationship is more complex when the community contains conspecific strains, or when mobile genetic elements are shared between species (**Fig. 1A**). In such cases, near-identical DNA sequences that belong to distinct strains are implicitly merged during the assembly process, resulting in partially overlapping configurations. To address this problem, we focus on finding clusters of contigs we call *anchors*, where (1) each anchor is a subset of the intersection of one or more overlapping configurations, and (2) no configuration contains contigs belonging to two distinct anchors. Anchor are operationally defined contig sets that provide a species-level representation of a potentially complex configuration space (**Supplementary Text**).

**Fig. 1.**
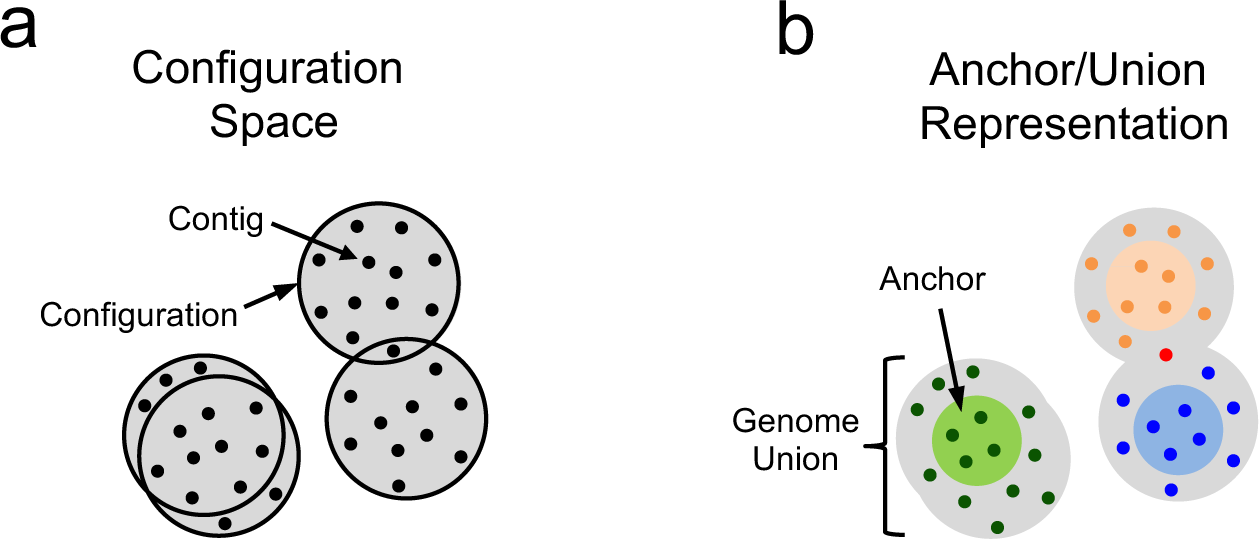
Genomic configuration space and an anchor-union representation. **(A)** Example with 4 configurations (large gray circles), each composed of contigs (black dots). Two related strains are represented by partially overlapping configurations. **(B)** Possible anchor-union pairs for the configurations in (A). There are 3 anchors (contigs within light-shade colored circles) and 3 matching genome unions, colored according to the anchor (dark shades). One contig is shared by two unions (colored red), representing a shared element, such as a plasmid. The two conspecific strains are represented by a single anchor-union pair.

To recover anchors from Hi-C contact maps we developed HPIPE, a probabilistic algorithm that explicitly addresses inter-cellular (spurious) contacts that confound the analysis of raw data. The algorithm infers a model that predicts the probability of an inter-cellular contact between two restriction fragments, as a function of fragment lengths and abundances (**Materials and Methods**). The model and anchors are co-optimized such that upon convergence each anchor is enriched for intra-anchor contacts relative to the model, and the contact enrichment between two different anchors matches the level predicted by the background model. In a final step, each anchor is extended into a *genome union*, by adding to it contigs that are enriched for anchor-specific contacts (**Supplementary Text**). A genome union (simply ‘genome’ throughout this work) represents the combined genome capacity of one or more conspecific strains that are associated with an anchor, potentially including shared genetic elements (**Fig. 1B**). The reduced representation of the genomic landscape using anchor-union pairs creates a unique opportunity to characterize genome structure in complex communities, which we exploited here to study HGT.

### Application of the method to the human gut

First, we tested our approach on two simple datasets. Application of the method to a simulated contact map generated for a community composed of 55 common gut bacteria, with varying degrees of relatedness and abundance (GOLD database^32^, **table S1**), resulted in 32 anchor-union pairs. Importantly, the probability of detecting a community member was associated with its abundance, confirming the non-biased nature of the method (**fig. S2**). Application of the method to published Hi-C data, generated from a synthetic microbial community composed of 5 strains^29^, resulted in the recovery of all species-level genomes, while merging two conspecific strains into a single anchor-union pair, confirming the ability of the method to work with real data (**fig. S3**).

We then applied the method to the contact map of subject A, resulting in 83 anchors (1.2Mb median anchor length). Thousands of spurious contacts between pairs of anchors were detected, yet the inferred background model was accurate in predicting this noise (Pearson=0.96, **Fig. 2A**). Each anchor was extended to a matching genome union, using stringent criteria (≥ 10-fold contact enrichment and ≥ 8 contacts, **fig. S4**). Contigs that were not associated with any anchor were discarded from downstream analyses. The resulting 83 genomes (2.7Mb median per genome) accounted for 75% of the estimated DNA mass in the sample, with preferential representation of the most abundant species (**Fig. 2B**).

**Fig. 2.**
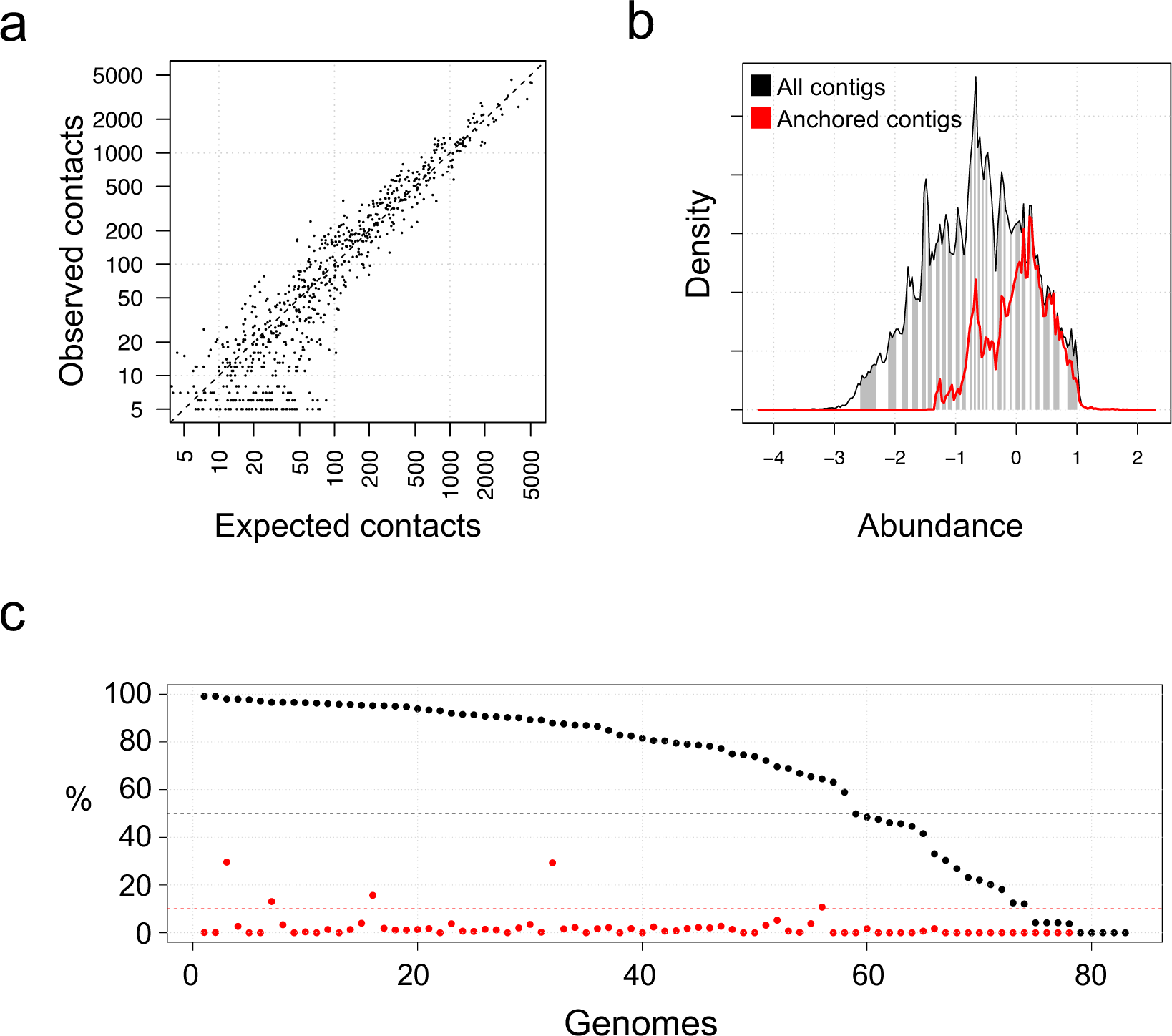
Genotyping complex microbial communities using Hi-C. **(A)** 83 anchor-union pairs were recovered for subject A. Shown is the expected number of inter-anchor spurious contacts (predicted by model, x-axis) vs. the observed number of inter-anchor contacts (y-axis). **(B)** A density plot of the relative abundance of all contigs from the metagenomic assembly (contigs >1k). The abundance (x-axis) is the enrichment of the contig read coverage over a uniform distribution of reads. The fraction of the assembly that was included in any recovered genome (‘anchored contigs’) is shown using a red line. White/gray stripes denote 10Mb bins. **(C)** Single-copy gene estimates of genome completeness percentage (in black) and contamination percentage (in red), and sorted according to completeness. Minimal completeness (50%) and maximal contamination (10%) thresholds depicted with dashed horizontal lines.

Genome completeness and contamination were estimated for all 83 genomes using the presence of universal single-copy genes^33^. Completeness was correlated with genome abundance (Spearman=0.36), and not with median contig length (representing assembly fragmentation, Spearman=-0.09), indicating that the major limiting factor for genome recovery in our community was sequencing depth. We examined 53 genomes that were draft-quality or better (>50% complete and <10% contaminated, **Fig. 2C**), and for each sought a single reference genome within the same species. We selected the most-closely related publicly-available genome, which was defined as the reference genome with the most conserved sequence (**Materials and Methods**). Nine of the 53 genomes lacked a species-level reference altogether, underscoring the still-incomplete characterization of the human gut microbiota, despite extensive study (**fig. S5**). Downstream analysis was limited to the remaining 44 genomes with a species-level reference.

Our results were comparable, in terms of genome number and quality, to a state-of-the-art metagenomic binning method^34^, and a recently published Hi-C binning method^35^ (**fig. S6**). However, the anchor-union approach we have implemented is unique in its ability to recover overlaps between genomes, making it ideal for studying within-host HGT, as we discuss next.

### Characterization of core and accessory genes

For each genome, we defined the *core genome* to be the portion of the genome with >90% nucleotide sequence identity to the reference, and the *accessory genome* to be the remaining portion of the genome (**Fig. 3**). We note that the use of only a single reference yields a conservative estimation of the accessory genome, since by definition cores diminish in size with the addition of strains to the analysis. Cores were on average 35% larger than their matching anchor, due to stringent anchor criteria (**fig. S7**). The accessory component was 25% (+/− 8.6%) of each genome, and accounted for 24,147 genes in total, grouped by synteny into 6391 accessory elements. Most cores showed high sequence conservation (>99%) with respect to their reference, while accessory components diverged by hundreds of genes, highlighting the contribution of HGT to strain diversification (**Fig. 4A**). We reasoned that if within-host HGT is ongoing in these subjects then it may be manifest by the sharing of mobile genetic elements between microbial hosts (i.e., donor and recipient strains). Indeed, a total of 264 elements (1086 genes) were robustly associated using Hi-C with multiple host genomes. Sharing was associated with genome sequence similarity but extended across family-level boundaries (**Fig. 4B**). The fraction of host pairs that shared elements increased from 4% to 84% as the host amino acid identity varied from 50% to 60%, confirming phylogenetic relatedness as a major determinant of HGT compatibility (**Fig. 4C**). Strikingly, 96 elements (307 genes) were shared by 3 or more microbial hosts, and some by as many as 6 hosts (**Fig. 4D**).

**Fig. 3.**
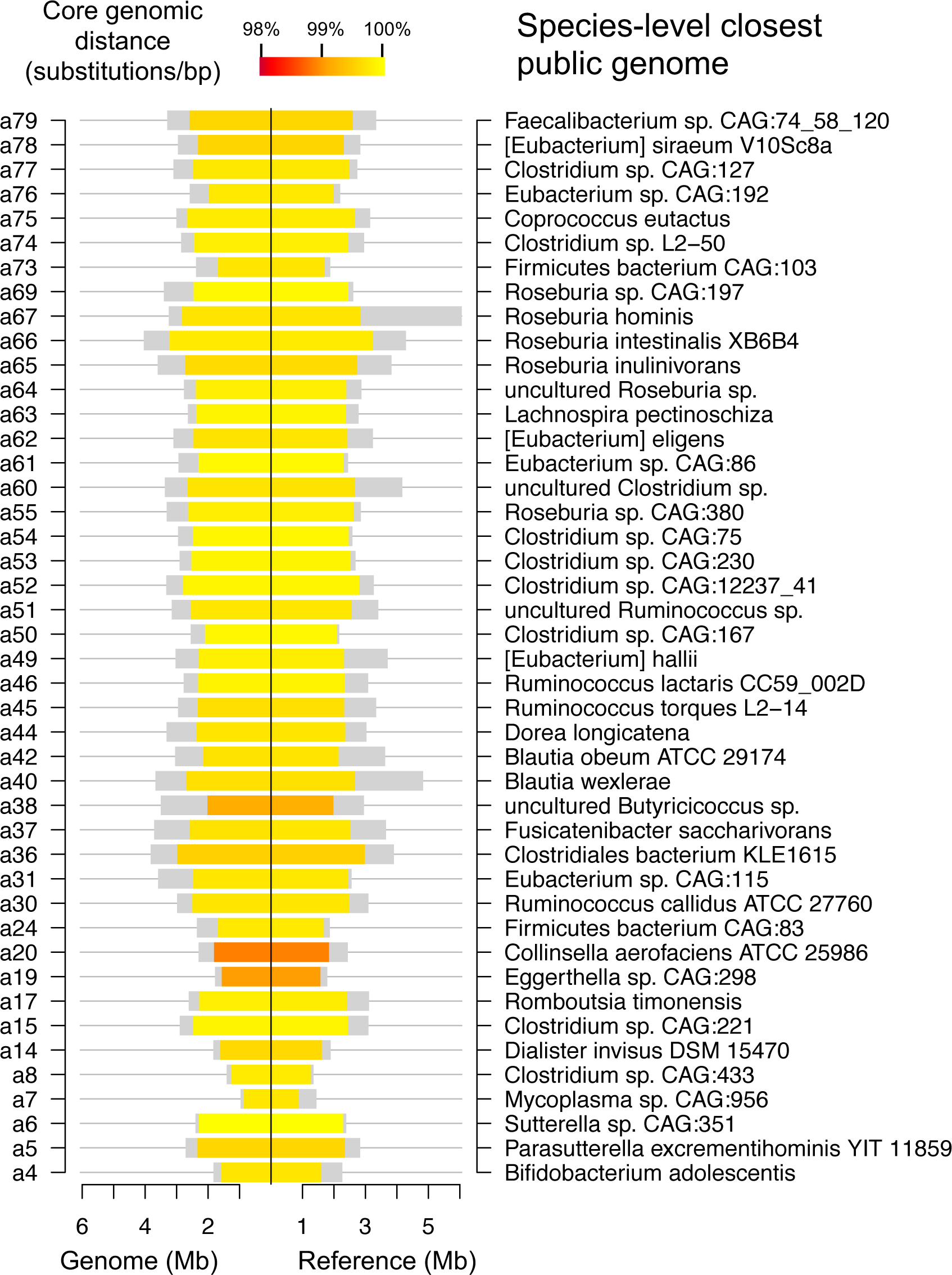
Determining cores and accessory genes. Core and accessory fractions for the 44 genomes that had a species-level reference. For both the recovered genomes (left) and the matching reference genomes (right), the core fraction is depicted using a colored rectangle, and the accessory fraction is depicted using a gray rectangle. Cores are colored according to the genomic distance (mean substitutions/bp) between cores and matching reference core.

**Fig. 4.**
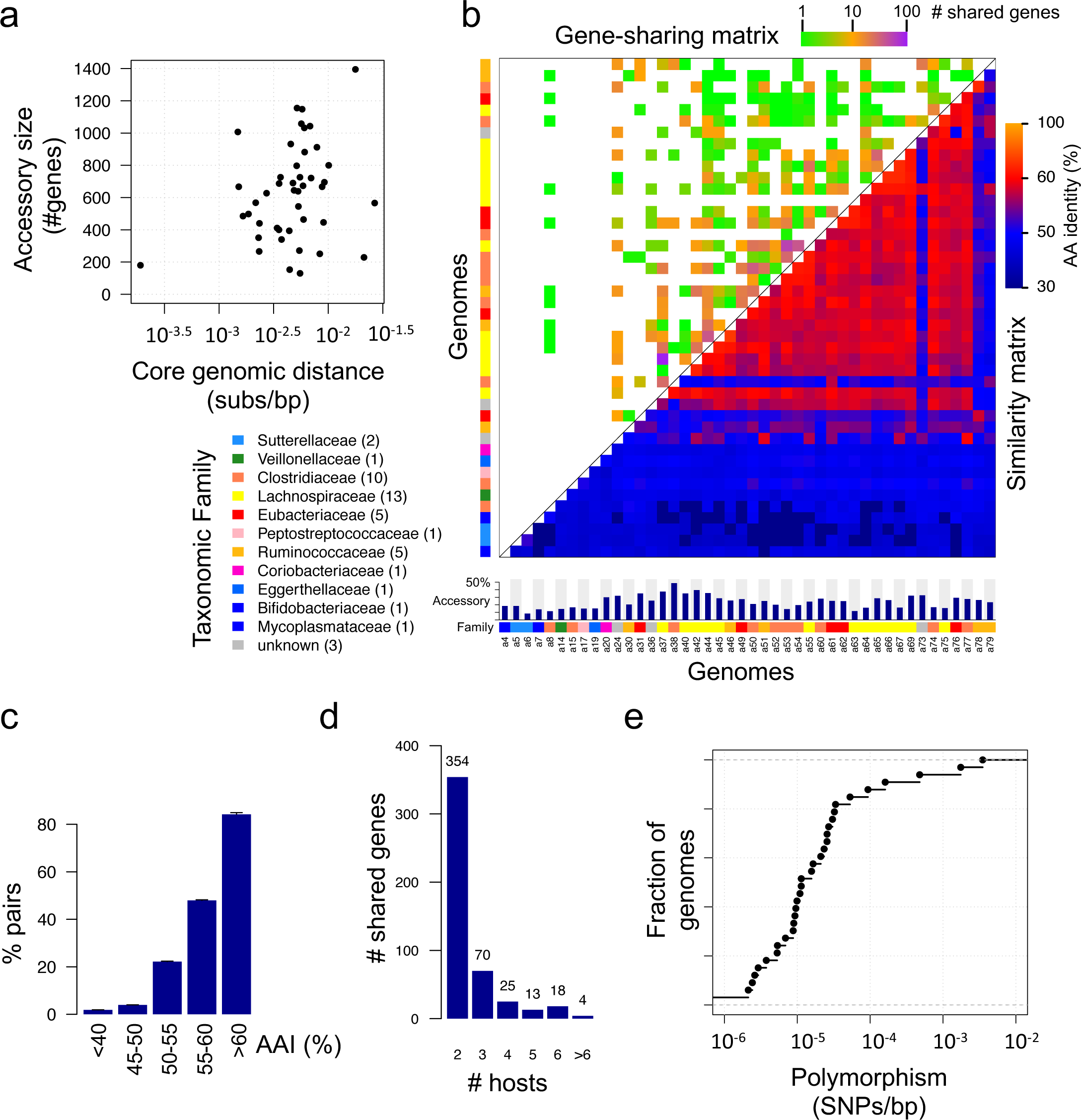
Attributes of accessory genes. **(A)** The substitution density within core genomes (x-axis) vs. the number of accessory genes (y-axis, genes that belonged to a recovered genome and were missing in the matching reference genome), for all 44 genomes that had a species-level reference. **(B)** Top left section of the matrix shows the number of shared genes and bottom right shows the mean amino acid identity (AAI). Genomes are sorted according to a hierarchical clustering based on AAI. Shown below the matrix is the size of the accessory fraction, and the Family taxonomic assignment for each genome (colored rectangles). The taxonomic family legend is shown with the number of genomes written in parenthesis. **(C)** The percentage of pairs of genomes that shared at least one gene, stratified by the sequence similarity (AAI) between the genome pair. **(D)** The number of shared genes, stratified according to the number of host genomes with which they were associated with. **(E)** The densities of intermediate SNPs (with allele frequency in the range 20-80%) within core genomes is plotted as an empirical distribution function, for 33 cores that had a read coverage of 10x or more.

To explore HGT dynamics and gut colonization history in greater depth, we estimated the within-host polymorphism levels of cores, by mapping metagenomic reads back onto the assembly and computing the densities of intermediate SNPs (single nucleotide polymorphisms with allele frequencies ranging from 20%-80%) (**Materials and Methods**). As shown in **Fig. 4E**, the majority of cores had low polymorphism levels (<10^−4^ SNPs/bp), consistent with a dominant clonal population that has experienced a recent within-host bottleneck (based on mutation accumulation rates in the range of 10^−8^ to 10^−5^ substitutions/bp per year, measured across diverse bacteria^36^). At the tail of the distribution, the most highly polymorphic cores likely represent distinct colonization events of conspecific strains, as they have polymorphism levels close to those that are typical for unrelated strains. Polymorphism levels were also estimated for 9 shared elements (out of 264), for which sufficient data were available (>10x coverage and >10kb in length). Strikingly, all 9 elements were highly clonal (<2*10^−4^ SNPs/bp), indicating they were likely spreading *in situ* (within the gut). To quantify HGT rates, we took a direct approach by using stool collected from the same person 10 years prior.

### Gut genome evolution over a 10-year period

We analyzed temporal changes in gene sequence and gene content, via metagenomic sequencing of a sample collected from the same subject 10 years prior to the genotyped sample. DNA was extracted and sequenced (320M reads), and reads were mapped to the 44 genomes described above. A single-nucleotide level investigation of mapped reads was able to differentiate between different scenarios (**Fig. 5A**). A total of 18 genome cores were not detected in the sample collected 10 years prior. The read coverage for 24 of the remaining 26 genomes was sufficiently high (>10x) to compute the core distance between the contemporary and past samples (**Fig. 5B**). A total of 3 strains accumulated low-level mutations (using a threshold of 10^−4^ substitutions/bp, based on empirical data^36^) and were classified as ‘persistent’, while the remaining 21 were classified as ‘replaced’.

**Fig. 5.**
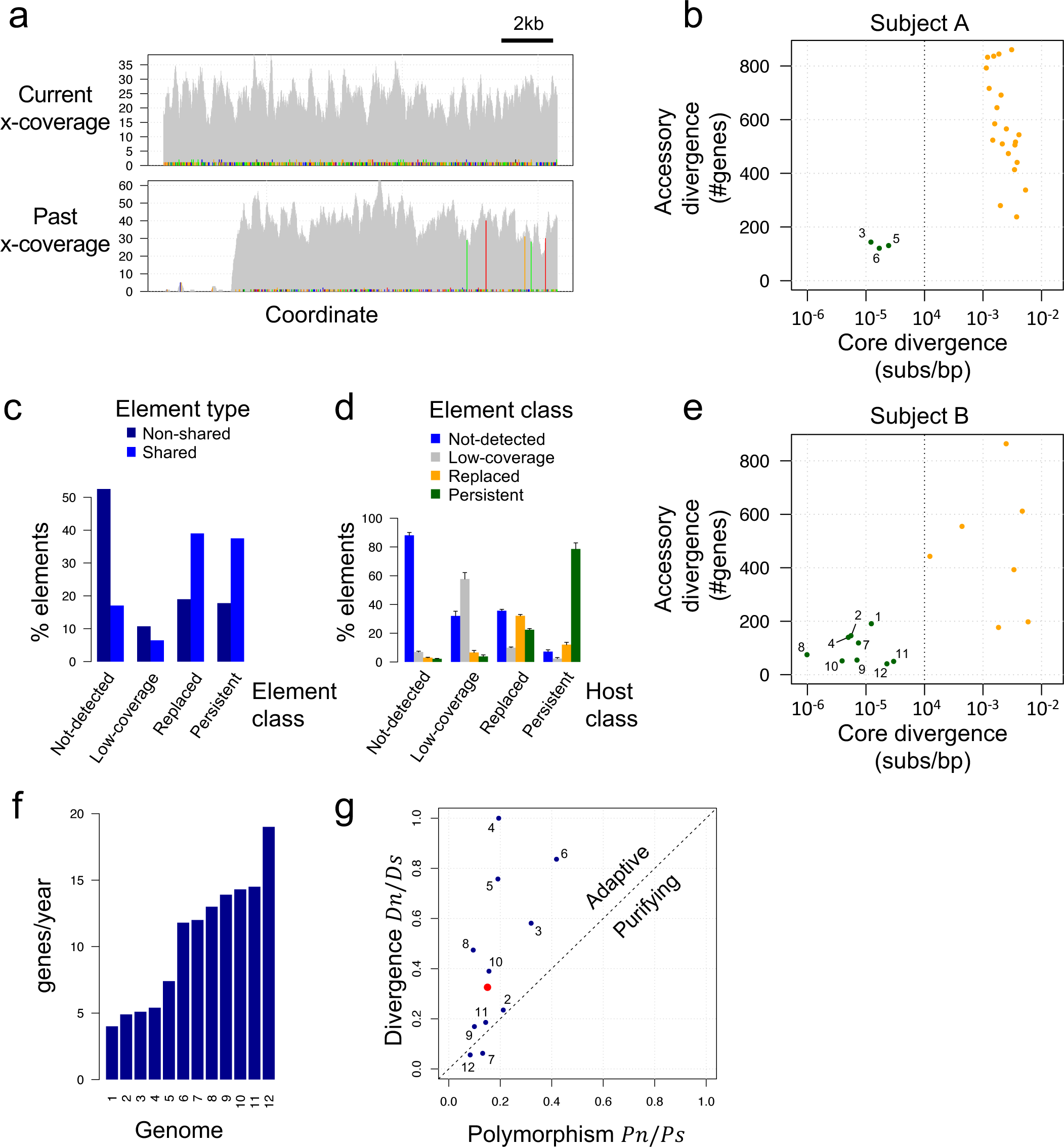
10-year community evolution. **(A)** Genetic changes along a 15kb segment (x-axis). Shown for the genotyped sample (top) and the sample collected from the same subject 10 years prior (bottom), is the number of read supporting each SNP (y-axis). SNPs that agree with the assembly are colored gray, and deviating SNPs are colored by nucleotide (A/C/G/T are colored red/blue/green/orange). Note in the 10-year profile the region on the left that has low read coverage (reflecting gene-content change), and the 5 divergent SNPs on the right (reflecting nucleotide-level changes). **(B)** Shown for 24 genomes that had >10x read coverage in the 10-year sample, is the core divergence (x-axis, substitutions/bp within cores) vs. the accessory divergence (y-axis, number of accessory genes classified as not-detected or replaced) over the 10-year period. Genomes are colored according to classification (persistent: green, replaced: orange), and the classification threshold (10^−4^) is depicted with a dashed vertical line. Persistent genome indices (as in Table 1) are numbered on the plot. **(C)** The distribution among element classes, stratified according to element type (shared and non-shared). Data is normalized so that each type sums to 100%. **(D)** The distribution among element classes, stratified according to host class. Data is normalized so that each host class sums to 100%. **(E)** Same panel B, for Subject B. **(F)** The gene turnover rate (y-axis) for the 12 strains classified as persistent, sorted according to the rate, with indices as in Table 1. **(G)** For the 12 persistent genomes, shown is the ratio between the density of synonymous (*PS*) and non-synonymous (*Pn*) polymorphic sites (x-axis), vs. the ratio between the density of synonymous (*DS*) and non-synonymous (*Dn*) divergent sites (y-axis). Persistent genome indices (as in Table 1) are shown. The divergence of genome #4 is plotted at 1 for visualization purposes, since no synonymous divergent sites were observed. Average values for all genomes (without genome #9, due to high levels of polymorphism) are plotted in red.

**Table 1.** Divergence summary for persistent genomes. Shown are all 12 genomes classified as persistent across both subjects. The SNPs/bp column shows polymorphic levels in the genotyped sample. Shown are the number of synonymous (#*Ps*) and non-synonymous (#*Pn*) sites polymorphic within the base sample, and the number of synonymous (#*Ds*) and non-synonymous (#*Dn*) sites divergent between the genotyped and the 10-year sample. Matching densities (*Ps,Pn,Ds,Dn*) were computed from raw count by normalizing for the total number sites of each type (synonymous and non-synonymous). P-values for the McDonald-Kreitman test were generated with the χ^2 test.

| Index | Subject | Genus | #genes | genes/year | SNPs/bp | #Pn | #Ps | #Dn | #Ds | Pn/Ps | Dn/Ds | P-value |
| --- | --- | --- | --- | --- | --- | --- | --- | --- | --- | --- | --- | --- |
| 1 | B | Unknown | 190 | 19 | 0 | 0 | 0 | 17 | 7 | - | 0.57 | - |
| 2 | B | Ruminococcus | 145 | 14.5 | 2.10E-05 | 18 | 20 | 5 | 5 | 0.212 | 0.235 | - |
| 3 | A | Clostridium | 143 | 14.3 | 1.65E-05 | 22 | 16 | 20 | 8 | 0.32 | 0.581 | 0.259 |
| 4 | B | Unknown | 139 | 13.9 | 1.39E-05 | 11 | 14 | 9 | 0 | 0.194 | >>1 | 0.003 |
| 5 | A | Clostridium | 130 | 13 | 9.54E-06 | 9 | 11 | 39 | 12 | 0.191 | 0.758 | 0.011 |
| 6 | A | Ruminococcus | 120 | 12 | 2.06E-05 | 25 | 14 | 25 | 7 | 0.418 | 0.837 | 0.198 |
| 7 | B | Faecalibacterium | 118 | 11.8 | 2.16E-05 | 10 | 19 | 2 | 8 | 0.131 | 0.062 | 0.392 |
| 8 | B | Bacteroides | 74 | 7.4 | 1.60E-05 | 14 | 35 | 2 | 1 | 0.095 | 0.474 | 0.165 |
| 9 | B | Alistipes | 54 | 5.4 | 0.002822 | 1134 | 2893 | 4 | 6 | 0.099 | 0.169 | 0.406 |
| 10 | B | Clostridium | 51 | 5.1 | 7.32E-06 | 6 | 9 | 5 | 3 | 0.156 | 0.39 | 0.304 |
| 11 | B | Lachnospira | 49 | 4.9 | 7.35E-05 | 68 | 113 | 32 | 41 | 0.143 | 0.186 | 0.355 |
| 12 | B | Faecalibacterium | 40 | 4 | 7.82E-05 | 28 | 82 | 6 | 26 | 0.083 | 0.056 | 0.434 |

We applied the same analysis to the 6391 accessory elements, classifying 3226 (51%) as ‘not-detected’, 1265 (19.8%) as ‘replaced’, and 1188 (18.6%) as ‘persistent’. The remaining 675 elements (10.6%) were detected 10 years prior but had low read coverage (<10x), confounding the differentiation between ‘replaced’ and ‘persistent’. Compared to elements associated with a single microbial host, shared elements were enriched for persistence and replacement (**Fig. 5C**). Analysis of element class, stratified by the associated host class, showed that elements did not always share the same history as their identified host (**Fig. 5D**). For example, out of 434 elements associated with persistent hosts, only 341 (78.6%) were classified as persistent, while 83 (19.1%) were classified as ‘not-detected’ or ‘replaced’, revealing extensive gene flux and recombination during that time period. Surprisingly, we also observed the reverse scenario, in which an accessory element seemingly predated its host in the gut: out of 2137 elements that were associated with ‘not-detected’ hosts, 45 (2.1%) were classified as ‘persistent’. These 45 elements provide direct evidence for dissemination of mobile elements within a single gut community, and a contrasting view to the idea of mobile elements as highly transient.

These intriguing findings led us to study a second individual (subject B), in an attempt to develop a more general understanding of HGT in the gut. In the case of subject B, we genotyped an early sample using Hi-C (650M Hi-C reads) and used a second sample collected 10 years later in order to track genetic changes (the reverse strategy to that used in subject A). The early sample of subject B generated 87 partial genomes, 44 of which were draft-quality or better and had a species-level reference (**fig. S8**). The genomes of subject B contained 25,327 accessory genes in total, grouped by synteny into 5860 elements; these genes accounted for 24% (+/− 10%) of each genome on average. DNA was extracted from the later sample of subject B and sequenced with 100M reads. Importantly, polymorphism levels and element classification distributions were remarkably similar between subjects (**fig. S9**). However, the gut community of subject B displayed greater levels of stability compared to subject A, with 9 bacterial hosts that were classified as persistent (**Fig. 5E**). By considering the 12 persistent strains identified in both subjects, we could estimate accessory gene turnover rates (**Table 1**). The rate of exchange of accessory genes among the persistent genomes was 4-19 genes/year (median 12 genes/year, **Fig. 5F**). These rates supersede by an order of magnitude previous estimates that were computed using long evolutionary branches^37^. These rapid HGT rates are in agreement with previous work that has shown that mutation accumulation rates are inversely correlated with the sampling time^36^.

To characterize whether selection was driving these rapid genetic changes, we performed the McDonald-Kreitman test^38^ for each genome, by comparing within-host polymorphism levels and divergence from the 10-year distant sample (shown in **Table 1**). The test indicated that some of the bacteria were evolving under strong adaptive (positive) selection during the 10-year period, while for others, the data were consistent with evolution in equilibrium (**Fig. 5G**). While the test was highly significant for only 2 genomes, pooling across all 12 genomes boosted the significance dramatically (𝜒^2^ test P<10^−7^). A systematic GO (gene ontology) analysis of the 152 core genes that contained non-synonymous substitutions identified GO categories that were enriched over a background composed of all predicted genes. Five categories were identified in both subjects, including signal transduction (hypergeometric test, P<0.0025) and nuclease activity (P<0.012). A matching analysis of the 1253 accessory genes that resided on elements putatively involved in within-host HGT (elements that were both classified as not-detected or replaced, and associated with a core classified as persistent), identified five enriched categories shared between the two subjects, including unidirectional conjugation (P<0.0012), DNA integration (P<0.01), and peptidoglycan catabolic process (P<0.02). The number of categories identified separately in the two subjects was significant for both core genes and accessory genes (𝜒^2^ test P<10^−16^), indicating that aspects of evolutionary processes were shared between the two subjects (see **table S2** for all identified categories). Together, the data suggested that gut bacteria evolve under a combination of varying levels of adaptive selection and extensive HGT.

### Specificity and prevalence of accessory genes in 218 individuals

To extend the results obtained from the 2 subjects and gain a population-based perspective on accessory genes, we used publicly available human gut metagenomes from 218 individuals (**table S3**). Reads were mapped using an efficient k-mer based approach to the assemblies from subjects A and B, and coverage vectors that spanned the 218 individuals were generated for all cores and elements (**Materials and Methods**). Each vector reflected the presence (>97% nucleotide identity) of either a core or an element across the cohort. The relationship between vectors of elements and of cores indicated the population-wide specificity of elements for their hosts, beyond the particular host-element associations observed in the genomes recovered from the two local subjects. At one extreme, a narrow-range element (for example, a species-specific bacteriophage) is expected to be present only when its host species is present, while at the other extreme, the presence of a broad-range element will be uncorrelated with the presence of the subject-specific host. Following this approach, we classified 12.4% and 15.1% of the elements of subjects A and B, respectively, as narrow-range, while the majority of the elements (69.6% for A and 73.8% for B) were classified as broad-range (**Fig. 6A**). When considering the contribution to any specific genome, broad-range elements accounted for an average of 12.3% and 13.8% of each genome of subject A and B, respectively, compared to only 3.5% and 4.7% for narrow-range elements (**Fig. 6B**). To obtain a more refined understanding of the host-specificity of broad-range elements we computed a specificity score, defined to be the Pearson correlation between the element vector and the vector of the host genome of that element in subjects A and B (or union of all host vectors, in the case of a shared element). Specificity scores ranged between 0 and 1, suggesting that a substantial portion of broad-range elements were decoupled from their locally inferred hosts (**Fig. 6C**). Unlike narrow-range elements which were rare, broad-range elements were found to be highly prevalent across the population, in levels comparable to microbial hosts (**Fig. 6D**).

**Fig. 6.**
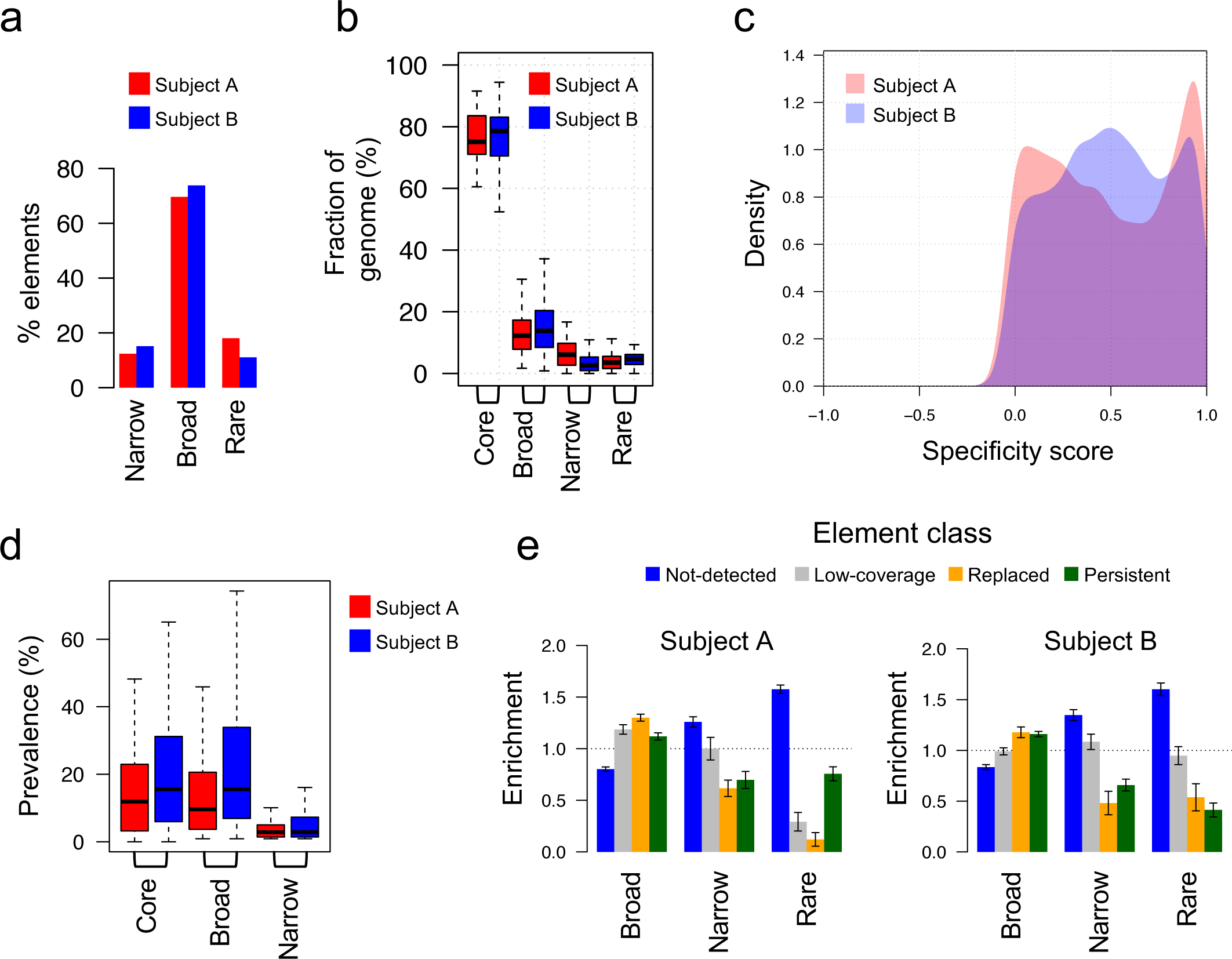
Population based perspective on accessory genes for the two subjects. **(A)** Elements were classified according to their distribution across 218 public gut metagenomic DNA libraries obtained from 218 individuals. The percentage of elements in each class for each of subjects A and B is shown. A ‘rare’ element was defined as an element detected in 0-2 individuals, and a ‘narrow-range’ element was defined as an element detected only in individuals in which one of its associated microbial hosts was also detected. All other elements were defined as ‘broad-range’. **(B)** Distribution across all 44 genomes of the genomic fraction (y-axis, percentage of genes out of the entire genome) of cores and broad/narrow/rare accessory fractions. **(C)** Population coverage vectors, spanning all 218 individuals, were computed for all accessory elements and cores. Shown is the density plot of element specificity scores, defined as the pearson coefficient between the vectors of broad-range elements and the vectors of their matching cores, colored by subjects. **(D)** The distribution of prevalence of cores, broad-range elements and narrow-range elements. **(E)** The enrichment of all combinations of population-based element classifications and evolution-based element classifications, over a null-model that assumes both classifications are independent.

We performed a GO analysis on these narrow- and broad-range elements, for both subjects (**table S4**). A total of 15 GO categories were enriched in narrow-range elements in both subjects, including viral capsid assembly (hypergeometric test P<0.0032), CRISPR maintenance (P<10^−8^), and cell motility (‘bacterial-type flagellum filament’, P<0.001). A total of 27 GO categories were enriched in broad-range elements in both subjects, including extrachromosomal circular DNA (P<10^−9^), unidirectional conjugation (P<10^−7^), DNA integration (P<10^−14^), transposition (P<10^−9^), DNA replication (P<0.0003), pathogenesis (P<0.004), virion assembly (P<0.005) and CRISPR maintenance (P<0.001). The overlap in terms of categories identified separately in the two subjects was significant (𝜒^2^ test P<10^−16^). We conclude that while both types of elements contain recombination genes and phage-related genes, broad-range elements stand out for their enrichment of conjugation genes, plasmid features, and pathogenesis-associated genes.

Finally, we compared turnover dynamics of the two types of elements, by tracking the evolutionary histories of narrow and broad elements over the ten-year period. Compared to narrow-range elements, broad-range elements were enriched for persistence and replacement (**Fig. 6E**). Together, the systematic analysis of hundreds of healthy individuals indicated that accessory elements are predominantly promiscuous and prevalent in human gut microbiotas.

## DISCUSSION

There is growing appreciation for the role of HGT in the evolution of adaptive traits in microbial communities, well beyond the roles described in earlier literature on the spread of virulence and antibiotic resistance. However, this understanding has arisen primarily from comparisons of distantly related strains available in public databases, which have been collected around the globe. Fundamental properties of HGT that emerge only in natural communities, including the extent, function and turnover rates of mobile elements, remain poorly understood. To address this problem, we developed a culture-free genotyping method to characterize genome dynamics in intact gut communities, resolving 88 strain-level genomes of gut bacteria from two subjects. Comparisons to publicly-available reference genomes suggested that accessory genes account for a quarter of each genome, on average. This striking gene-content variation can be attributed to a combination of gene gains (via HGT) and gene deletions. Temporal analysis over a 10-year period revealed complex dynamics, including colonization/extinction events, strain replacements, and importantly, *in situ* evolution of persistent strains. The presence of persistent strains allowed us to make a direct estimate of HGT rates, and provided evidence for adaptive evolution in some of the core genomes. Finally, a population-based analysis indicated that the accessory genome is dominated by broad-range elements that are prevalent in human gut microbiotas and have varying degrees of specificity for the host genome in which they were identified.

The genotyping approach presented here combines Hi-C with a probabilistic framework and uses anchor-union pairs to represent complex population structures. The approach is well poised to make significant inroads towards an understanding of complex microbial community structures and dynamics, such as those found in soil, which routinely defy standard binning and other approaches. While promising alternative approaches based on long-reads exist^39, 40^, Hi-C is notable for its ability to provide proximity information across millions of base-pairs of contiguous sequence, including inter-molecular contacts, as demonstrated by the association of plasmids with their respective host chromosomes. The limitations of the method include possible strain interference (i.e., fragmented assemblies due to the presence of conspecific strains) and possible differing experimental efficiencies (e.g., differential lysis of cell walls or resistance to restriction enzymes). However, a more obvious limiting factor is sequencing depth; a back-of-the-envelope calculation suggests that the allocation of 1 billion reads results in an abundance detection limit of 0.1%, and the detection limit is expected to drop linearly with sequencing depth.

Recent attention to microbial *in situ* evolution, long appreciated as a primary ecological process underpinning community assembly and diversification, has provided an unprecedented view on genome dynamics in natural environments, in real time, and with implications for human health. Other recent work provides independent evidence for HGT and adaptive evolution in the human gut, using an isolate-based approach focused on *Bacteroides fragilis*^41^ and a reference-based approach using the pangenomes of 30 common gut species^42^. The culture-independent and reference-free approach presented here opens the door to studying fundamental aspects of microbial evolution in complex and poorly characterized environments.

## ACKNOWLEDGMENTS

We thank Megan Kennedy for help in processing clinical samples and the members of the Relman and Holmes laboratories for invaluable discussion and feedback.

### Funding

This work was supported by NIH R01AI112401 (D.A.R. and Susan Holmes), EMBO Long-Term Fellowship ALTF 772-2014 (E.Y.), and the Thomas C. and Joan M. Merigan Endowment at Stanford University (D.A.R.).

### Competing interests

None declared.

### Author Contributions

E.Y. and D.A.R. designed the study. E.Y. developed the methodology and performed the analysis. E.Y. and D.A.R. reviewed the analysis and wrote the manuscript.

### Data and materials availability

Unprocessed, raw DNA sequence reads are available in the SRA database, under project PRJNA505354. HPIPE is available for download as an open-source tool at https://github.com/eitanyaffe/hpipe.

## MATERIALS AND METHODS

### Sample collection and shotgun procedure

Subjects A and B are healthy Western adult males who have not used antibiotics for at least 6 months prior to sampling. Fresh stool was collected and stored at −80C until processing. To generate standard DNA libraries (for the metagenomic assembly and for the temporal comparison), DNA was extracted using the AllPrep DNA/RNA Mini Kit (Qiagen), sheared and size-selected (>300bp), and paired-end sequenced using Illumina HiSeq 2500.

### Hi-C procedure

To generate the Hi-C DNA libraries, 50-100mg of stool was suspended in 10ml cold PBS, vortexed for 20min at RT, and spun down at 20g for 10m at 4C. The supernatant was centrifuged at 5000g for 10min, the resulting pellet was washed 2 more times in cold PBS, and the final microbial pellet weight **W** (in mg) was recorded. The pellet was suspended in 5.5ml PBS, fixated with 2.5ml formaldehyde 16% (final 5%) for 30min at RT and 30m on ice. The reaction was quenched with 1525ul glycine 2.5M (final 0.4M) for 5min at RT and 15min on ice. Fixated cells were washed twice with 10ml cold PBS, suspended with 4x**W**ul of H_2_O (4 times the recorded microbial pellet weight **W**), and 50ul aliquots of the fixated cell pellet were stored at −80C. For lysis, 10ul fixated input (∼2mg of microbial pellet) were suspended in 190ul TE and 1.1ul Ready-Lysozyme 36KU/ul (final 200U/ul), and incubated 15min at RT with occasional pipetting. Next, 10ul SDS 10% (final 0.5%) was added and samples were incubated for 10min at RT (total reaction volume, 200ul). For digestion, 150ul H_2_O, 50ul 10x DpnII buffer, 50ul Triton 10% (final 1%), and 50ul DpnII restriction enzyme (final 5U/ul) were added, and samples were incubated at 37C for 3hrs (final reaction volume, 500ul). Samples were incubated 10min with 25ul SDS 10% (final 0.5%) at RT. For ligation, 800ul Triton 10% (final 1%), 800ul 10x T4 buffer, 80ul 10 mg/ml BSA and 5800ul H_2_O and 20ul T4 ligase (final 2000U/ul) were added, and the sample was incubated for 4 hours at 16C (final reaction volume, 8ml). Following ligation, 100ul Proteinase K 20ug/ul (final 250ug/ml) was added and samples were incubated overnight at 65C. DNA was then cleaned with phenol-chloroform, precipitated in ethanol, suspended in 500ul TE, transferred to 1.5ml tubes, and incubated 1hr at 37C with RNase 0.5ug/ul (final 30ug/ml). DNA was cleaned with 2 more rounds of phenol-chloroform, ethanol precipitated, washed twice with 70% ethanol, and eluted in TE. DNA was sonicated, size-selecting for fragments 500-800bp and paired-end sequenced using Illumina HiSeq 2500.

### Preprocessing raw reads

Identical duplicate reads were removed, reads were quality-trimmed using Sickle1 with default parameters, adaptor sequences were removed using SeqPrep2 (min length of 60nt), and human sequences were removed using DeconSeq^3^ (alignment coverage threshold 10%, identity threshold 80%), resulting in unique high-quality non-human paired reads.

### Metagenomic assembly

*De novo* metagenome assembly was performed using MEGAHIT^4^ with parameters “--min-contig-len 300 --k-min 21 --k-max 141 --k-step 12 --merge-level 20,0.95”, and filtering out contigs shorter than 1kb. For mapping reads onto the assembly, the first 10nt of each read were trimmed, and the following 40nt were mapped using BWA-MEM^5^ with default parameters. Low quality or non-unique reads (>0 mismatches, <30nt match length or mapping score <30) were filtered out.

### Hi-C contacts

Contigs were pairwise aligned using Mummer^6^, identifying identical stretches of sequence (>=20nt long) shared between pairs of contigs. If the two sides of an inter-contig Hi-C paired read mapped up to 2000bp away from a perfect alignment region, the read was filtered out. The restriction enzyme that was used (DpnII) induces a partitioning of all contigs into restriction fragments. Every Hi-C ligation event (‘contact’) occurs between two fragment ends. To infer a contact from a mapped read pair, the contig was scanned from the mapped read coordinate, in the direction of the mapped read strand, until the first DpnII restriction site was reached, separately for both sides of each read pair. To minimize sequencing amplification noise, contact multiplicity was ignored, i.e. only unique contacts were considered.

### Inference of anchor-union pairs

We defined the *abundance* of a contig c to be the normalized read-coverage 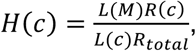 where *R*(*c*) is the number of Hi-C reads that mapped to *c*, *R_total_* is the total number of reads in the library, *M* is the set of all contigs in the metagenome assembly, and *L*(*X*) is the total length in base pairs of a contig set *X* ⊆ *M*. We defined the *weighted mean abundance* of a contig set *C* ⊆ *M* to be 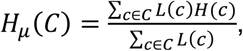 the *weighted standard deviation* to be 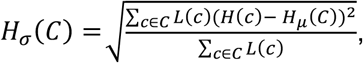 and the *abundance z-score* of a contig *c* ∈ *C* to be 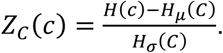 We modelled the probability of a spurious contact between two fragment ends *x*, *y* as:

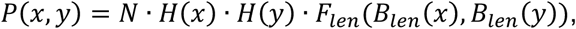

where N is a normalizing constant, *H*(*x*) and *H*(*y*) are the abundances of the contigs on which the fragments with ends *x* and *y* reside (respectively), and *F*_len_ is a function that transforms a pair of binned values *B_len_*(*x*), *B_len_*(*y*) of fragment lengths into a single empirical correction factor.

Given a spurious model *P* and constants *α*, *β* ∈ ℝ, we denoted two disjoint contig sets *X*, *Y* ⊆ *M* as *(α*, *β)-associated* if (1) *X* and *Y* were connected by at least *α* contacts, (2) the number of connecting contacts was at least *β*-fold enriched over the spurious contacts predicted by the model *P*, and (3) the false positive binomial probability for the observed contacts was below 10^−6^. The inferred *anchors* were a disjoint collection of contig sets 𝔸, for which each anchor *A* ∈ 𝔸 satisfied these five conditions:

(A1) **Clique**: Over 90% of pairs of contigs *a*, *b* ∈ *A* were associated by one or more contacts.

(A2) **Association**: Every contig *a* ∈ *A* and *A*\*a* were associated, with *α* = 5, *β* = 1.6.

(A3) **Uniqueness**: No contig *a* ∈ *A* and *A*′ ∈ 𝔸\*A* were associated, with *α* = 5, *β* = 1.6.

(A4) **Size**: Every contig *a* ∈ *A* was ≥10kb, and the total length of contigs in *A* was ≥200kb.

(A5) **Abundance**: *H_σ_*(*A*) ≤ 0.2, and for all *a* ∈ *A* the z-score *Z_A_*(*a*) ≤ 1.5.

The model *P* and anchors 𝔸 were inferred simultaneously. Briefly, seed anchors were computed using hierarchical clustering. A seed model was inferred over the seed anchors using maximum likelihood. Contigs that were associated with multiple anchors were discarded sequentially until convergence. Finally, anchors that were small or had a large abundance variance were discarded.

The matching genome union *A* ⊆ *G* was generated by including any contig *c* ∈ *M* that satisfied:

(G1) **Association**: The contig *c* and *A*\*c* were associated, with *α* = 8, *β* = 10.

(G2) **Anchor support**: The association was supported by at least 2 anchor contigs.

(G3) **Contig suppor**t: The association was supported by least 50% of the fragment ends within the contig *c*.

Default HPIPE parameters were tuned to favor precision over sensitivity, and are customizable. See the **SI methods section** for a complete description of the algorithm.

### Validation on simulated communities

Reference genomes for 55 common gut bacteria (GOLD database^7^) were downloaded from NCBI (**table S1**). The contigs of each reference genome were concatenated into a single circular pseudo contig. Genomes were ordered randomly and assigned an x fold-coverage value that ranged between 1 and 1000 following a geometric progression. To generate the assembly library, random read pairs (2×150bp) were generated, given the assigned x-coverage for all genomes, resulting in a total of 120M read pairs. The distribution of the distances between read pairs was a Gaussian with an offset: 200bp + N (mean=800bp, sd=200bp). To generate the Hi-C library, 100M random read pairs were generated as follows. A total of 1% of reads were allocated to be spurious reads, and were associated with two independently selected genome coordinates chosen according to genome abundance. The remaining 99% reads were assigned to genomes according to their abundance. Within each genome the distance between 50% of reads was uniformly distributed and the distance between the remaining 50% was distributed following a power law with an exponent of −1. HPIPE was run on the assembly and Hi-C library using default parameters.

### Validation on synthetic community

Raw Hi-C sequencing data were downloaded for a clonal synthetic community^8^, which was composed of 5 microbial strains: *Pediococcus pentosaceus* (ATCC 25745), *Lactobacillus brevis* (ATCC 367), *Burkholderia thailandensis* (E264) and two strains of *Escherichia coli* (BL21 and K-12). Matching reference genomes were downloaded from NCBI. An assembly library with an x-coverage of 100 was simulated, as described for the simulated community above. HPIPE was run on the simulated assembly library and downloaded Hi-C data using default parameters.

### Comparison to alternative methods

MetaBAT2 (version 2.12.1) was applied to the metagenomic assembly and the supporting reads of the assembly of Subject A, using default parameters. Bin3C (downloaded from GitHub on March 2019) was run on the metagenomic assembly and the raw Hi-C DNA library of Subject A, following the guidelines supplied by the bin3C authors, and using default parameters.

### Genome sequence similarity

Genes were predicted on all contigs using MetaGeneMark^9^ and were self-aligned using DIAMOND^10^ (sensitive mode, E<0.001). For all pairs of genome unions, if there were at least 12 aligned gene pairs (>30% identity and >70% coverage), the average amino acid identity (AAI) was computed by averaging the alignment identities (correcting identity for partial gene coverage, to reflect alignment over all of the gene), and otherwise it was set to 0. To generate the sequence similarity matrix (**Fig. 4B**), genome unions were clustered using hierarchical clustering, using AAI as the similarity metric and merging clusters using the ‘average’ method.

### Taxonomic affiliation

Single-copy gene analysis was performed using CheckM^11^. Genomes which were less than 50% complete or more than 10% contaminated were discarded from downstream analysis. Predicted genes were blast-aligned to UniRef100 (Downloaded in December 2015) using DIAMOND (sensitive mode, E<0.001). For each genome union, UniRef homolog genes (>30% identity and >70% coverage) were converted into one or more corresponding NCBI taxonomic Entrez entries, and organized on a taxon tree. The number of homolog genes was propagated up the tree. A *species taxon* was determined to be the species-level tree node that (1) had the maximal gene count among all species-level nodes, and (2) had one or more available reference genomes in the GenBank database^12^ (Downloaded in May 2018).

### Species-level reference genomes

For each genome union, all reference genomes of the species taxon, as defined by the GenBank database, were downloaded from NCBI. For every candidate reference genome, a bi-directional mapping was performed by splitting the genome union and the reference genome into overlapping 100bp windows (sliding 1bp along the genome), and mapping in both directions using BWA-MEM^5^ with default parameters. For both the genome union and the reference genome, each coordinate was assigned the maximal sequence identity of all windows that contained it, producing an *identity track* for both directions of mapping. The *alignable fraction* was defined as the portion of the genome union that was successfully mapped, averaged over both directions of mapping. The *nearest reference genome* was selected to be the reference genome for which the alignable fraction was maximal.

### Core and accessory fractions

For each genome that had a nearest reference genome, a gene-level nucleotide identity vector, was computed by averaging the mapping identity over entire genes. Genes for which the identity was 90% or more were defined as core genes, and the remaining were defined as accessory genes. A genome was classified as ‘no-reference’ if (1) there the assigned species taxon had no reference genomes in GenBank, or (2) the fraction of core genes was <50%. This resulted in 9 putative novel genomes for subject A and 13 putative novel genomes for subject B. Accessory genes were grouped into accessory elements according to synteny, i.e. if they appeared sequentially within a contig. Elements for which the gene x-coverage z-score distribution had a high standard deviation (>4) were removed from downstream analysis (in total <2.5% of elements were removed in this manner).

### Polymorphism levels

Complete assembly read sides were mapped onto the assembly using BWA-MEM^5^ with default parameters. Only matches that were 100bp or more, with a maximal edit distance of 2 and a score of 30 were used. A nucleotide-level vector with the allele frequency for all 4 nucleotides was computed by parsing the SAM alignment result. A nucleotide coordinate was called *intermediate* if (1) the allele frequency f satisfied 20%<f<80%, and (2) there were at least 3 supporting reads for the allele. The *polymorphism level* (i.e. the standing variation) for a gene-set (core or element gene-set), which had a sufficient read coverage (>10x), was defined to be the mean density of intermediate SNPs over the gene-set, discarding a 200bp margin near contig edges.

### 10-year core and element classification

The secondary sample, taken 10 years apart, was mapped onto the assembly using BWA-MEM, and generating a nucleotide-level vector with the allele frequencies as for the standing variation. A nucleotide coordinate was called *fixed* if (1) the dominant nucleotide was different from the assembly reference nucleotide, (2) the allele frequency was at least 95%, and (3) there were at least 3 supporting reads for the allele. The *substitution density* for a gene-set (core or element gene-set), was defined to be the density of fixed coordinates over the gene-set, discarding a 200bp margin near contig edges. A gene-set (core or element) was classified as *detected* if >90% of the genes had a median read coverage of 1x or more, and it was classified as *not-detected* otherwise. A detected gene-set was further classified as *high-detected* if (1) the median read coverage over the entire gene-set was at least 10x, and was classified as *low-coverage* otherwise. High-detected gene-sets were further classified as *persistent* if the substitution density over the gene-set was <D_t_, and classified as *replaced* otherwise. The threshold D_t_ was set to 10^−4^, based on empirical estimates of mutation accumulation rates in bacteria, that range between 10^−8^ and 10^−5^ substitutions/bp per year^13^. The *accessory divergence* of a genome was the total number of accessory genes associated with the genome that were on elements classified as not-detected or replaced.

### McDonald-Kreitman test

Test values were computed for each of the 12 genomes that were classified as persistent across both subjects. Synonymous and non-synonymous sites were determined using Translation Table 11 (NCBI). The number of synonymous (#*PS*) and non-synonymous (#*Pn*) polymorphic sites were computed per core using intermediate SNPs. The number of synonymous (#*DS*) and non-synonymous (#*Dn*) divergent sites were computed per core using fixed SNPs. Matching densities (*PS*, *Pn*, *DS*, *Dn*) were computed from raw count by normalizing for the total number sites of each type (synonymous and non-synonymous). P-values for the McDonald-Kreitman test were generated using the 𝜒^n^ test over (#*PS*, #*Pn*, #*DS*, #*Dn*).

### Gene ontology enrichments

Enrichments for GO (Gene ontology) categories were computed as follows: All Uniref100 hits were transformed into GO categories, using the Uniparc and Uniprot databases as intermediates. To generate the p-values reported for a given GO category and a selected set of predicted genes, a hypergeometric test was performed by comparing the selected set to a background set composed of all predicted genes.

### Population presence analysis

218 human gut metagenomic DNA libraries collected from distinct subjects were downloaded from the HMP and the EMBL-EBI repositories (**table S4**). Each of the 218 subject libraries was converted to a *k-table* (k=16), by counting the frequency of all k-mers across the library reads. The following analysis was performed separately for subjects A and B. Each k-table was projected on each predicted gene, generating a 1-bp vector of k-mer frequencies. The *gene coverage* was defined as the median k-mer frequency over the entire gene vector. The *gene fraction* was defined as the fraction of the gene vector that was covered by segments of hits that were at least *q* = 30 long. The value of the parameter *q* was selected to balance between false positives and the detection limit, that was estimated to be 96.66% (100 − 100⁄*q*), assuming substitutions are disturbed uniformly. A gene *g* was called *present* in the library of subject *i* if (1) the gene fraction in library *i* was at least 80%, and (2) the gene coverage in the library was at least 2. The *presence value* 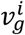 was set to be the gene coverage if the gene was called as present in the subject library, and set to zero otherwise, resulting for each gene *g* in a *gene presence vector* 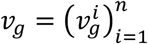 that spanned all 218 subjects.

For a gene-set *x* (either a core or an element), the *set presence vector v*_*x*_ was defined to be a per-coordinate median over the presence vector of the genes in the gene set: 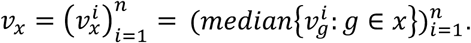 In this manner presence vectors for all elements and their associated cores were computed. The *detected subject set S*(*v*) of a presence vector *v* was defined to be 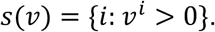 For each element *e* and its matching set of host cores *H_e_* (one or more hosts), the *element host presence vector* was defined to be 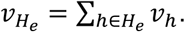 The element was classified as *rare* if the detected subject set 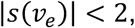 as *narrow* if 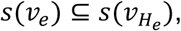 and as *broad* otherwise. The *element-host specificity score* was defined to be the Pearson correlation between the presence vectors 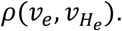

### Association between 10-year classification and population classification

Each element was classified into 3 classes using the 10-year dataset (not-detected/low-coverage/replaced/persistent), and into 3 classes using the population dataset (rare/broad-range/narrow-range). The observed number of elements classified under all 12 combinations of classification pairs was counted. To generate **Fig. 6E** the observed number of elements was compared to the expected number of elements was estimated using a generalized Bernoulli distribution.

## 1 Basic definitions

### 1.1 Contig abundance

We defined the *abundance* of a contig c to be the normalized read-coverage 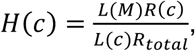 where *R*(*c*) is the number of Hi-C reads that mapped to *c*, *R_total_* is the total number of reads in the library, *M* is the set of all contigs in the metagenome assembly, and *L*(*X*) is the total length in base pairs of a contig set *X* ⊆ *M*. We defined the *weighted mean abundance* of a contig set *C* ⊆ *M* to be 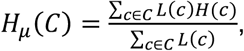 the *weighted standard deviation* to be 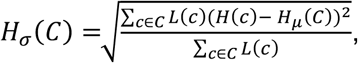 and the *abundance z-score* of a contig *c* ∈ *C* to be 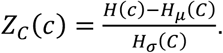

### 1.2 Genome configurations and linkage

We use the term *genome configuration* to refer to a set of contigs that represent the genomic capacity (including extra-chromosomal DNA) of a clonal population of cells in the community. We call a pair of contigs *linked* if there is one or more configuration that contains both contigs, e.g. they are both a part of the same genome. We call any Hi-C contact that associates two non-linked contigs a *spurious* contact, since it is a result of experimental noise (likely due to an inter-cellular ligation event).

### 1.3 Spurious contact model

We represent a Hi-C contact map as an indicator function *I*(*x*, *y*), that equals 1 if there is one or more contacts associating the ends *x*, *y* from two fragments, and equals 0 otherwise. We model the probability of a spurious contact between two fragment ends *x*, *y* as:

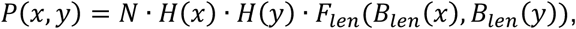

where *H*(*x*) and *H*(*y*) are the abundances of the contigs on which the fragments with ends *x* and *y* reside (respectively), *F_len_* is a function that transforms a pair of binned values *B_len_* (*x*), *B_len_* (*y*) of fragment lengths into a single empirical correction factor, and *N* is a normalizing constant.

For two contig sets *X*, *Y* we define the *observed contacts* 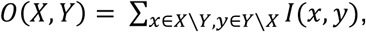 the *expected contacts* 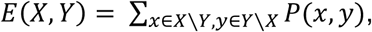 and the *contact enrichment score* 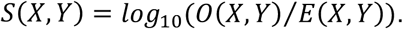 The distribution of the observed number of contacts between two non-linked contig sets *X*, *Y* is approximated using a binomial distribution. The *false positive probability Q*(*X*, *Y*) is the probability of observing at least *O*(*X*, *Y*) contacts, assuming a binomial distribution of contacts 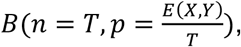 where *T* is the total number of observed contacts.

### 1.4 Contig graph

We define a *contig graph* over the contigs, where each contig is a vertex and each pair of contigs that is associated by a contact as connected by an edge. *N*(*c*) denotes the neighbors of a contig *c* in the contig graph. The *shared neighbors metric D* between two contigs *c*_1_, *c*_1_ is defined to be *D*(*c*_1_, *c*_1_) = |*N*(*c*_1_) ∩ *N*(*c*_2_)|. The *clique degree* of a contig set *C* is defined to be 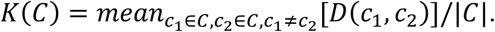 Note that if *C* is a clique (i.e., there are contacts between all pairs of contigs in *C*) then *K*(*C*) ≥ 1.

## 2 Genome anchors and genome unions

### 2.1 Genome anchors

A disjoint collection 𝔸 of contig sets 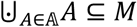 is called an *anchor collection* if it satisfies these five conditions:

(A1) **Clique**: Threshold on the clique degree of the anchor.

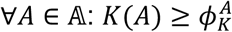

(A2) **Association**: Each contig of an anchor must be associated with the anchor.

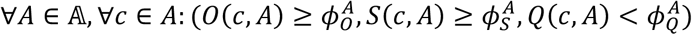

(A3) **Uniqueness**: Each contig of an anchor must not be associated with any other anchor.

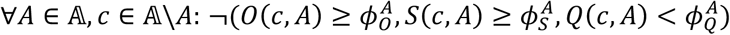

(A4) **Size**: Thresholds on contig and anchor length (in basepairs).

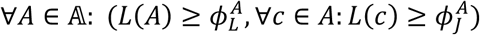

(A5) **Abundance**: Thresholds on the standard deviation and z-scores of the abundance.

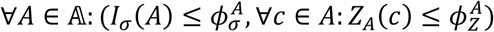

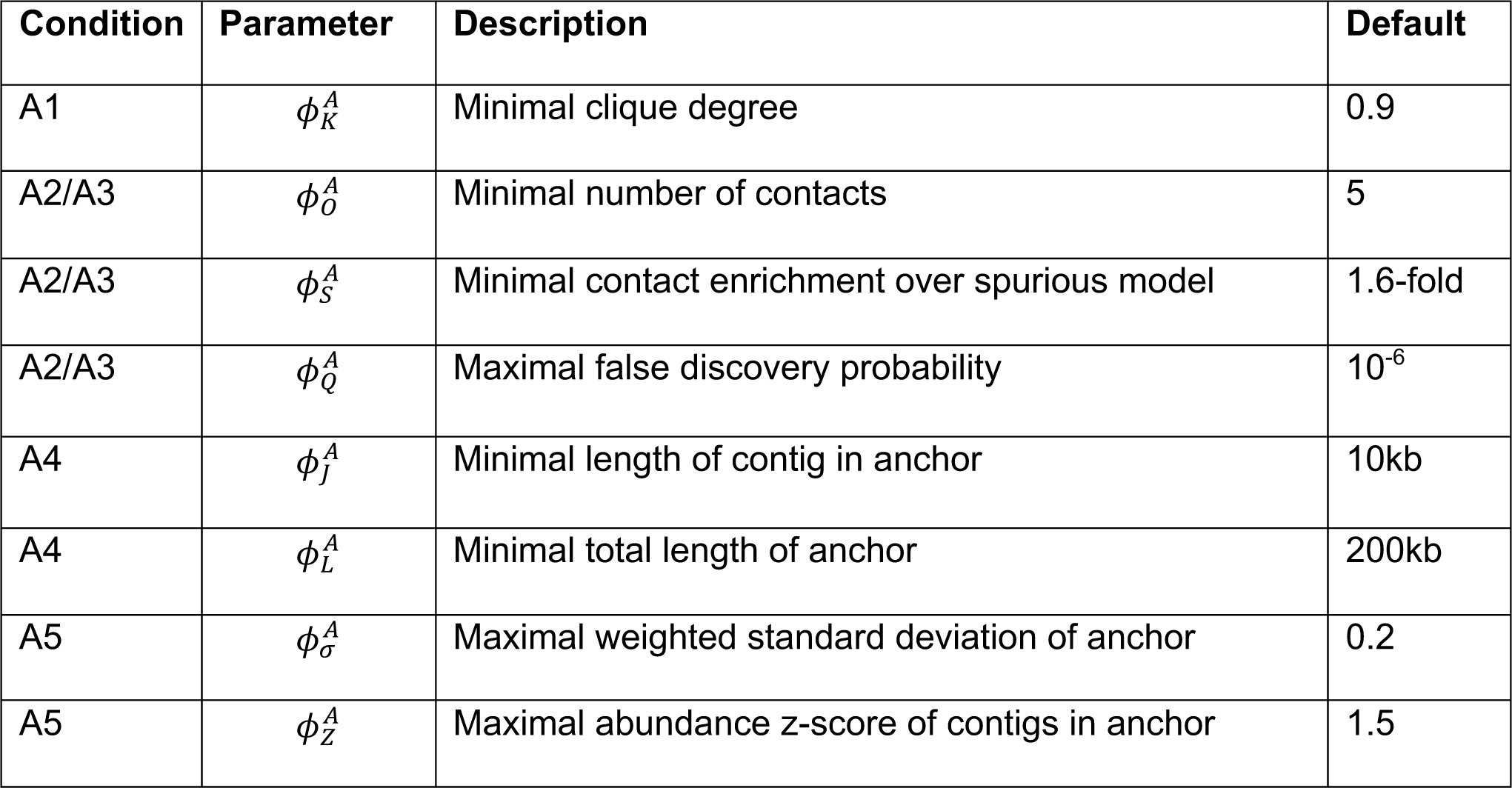

Default parameters were selected to favor precision over sensitivity, and are customizable.

### 2.2 Genome unions

Given an anchor collection 𝔸, each anchor *A* ∈ 𝔸 is extended into a *genome union* by including all contigs that are associated with the anchor, according to the Hi-C contact map. Taking a stringent approach, we filter out contig-anchor pairs for which the contacts are limited to a small portion of either the contig or the anchor. We break down each contig *c* into 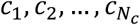 bins of fragment ends, where *N_c_* = *min* (10, *f*(*c*)), and *f*(*c*) is the number of fragments on *c*. We define the *contig support* of the pair (*c*, *A*) to be 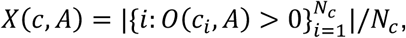 and the *anchor support* of the pair to be *Y*(*c*, *A*) = |{*a* ∈ *A*: *O*(*c*, *a*) > 0}|. The *genome union G*(*A*) is then defined to be all contigs *c* ∈ *M* that satisfy these three conditions:

(G1) **Association**: The contig must be associated with the anchor.

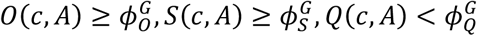

(G2) **Anchor support**: Threshold on the anchor support of the association.

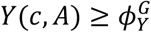

(G3) **Contig support**: Threshold on the contig support of the association.

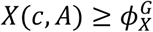

| Condition | Parameter | Description | Default |
| --- | --- | --- | --- |
| G1 | $\phi_O^G$ | Minimal number of contig-anchor contacts | 8 |
| G1 | $\phi_S^G$ | Minimal contig-anchor contact enrichment over spurious model | 10-fold |
| G1 | $\phi_Q^G$ | Maximal false discovery probability | $10^{-6}$ |
| G2 | $\phi_Y^G$ | Minimal number of supporting contigs in anchor | 2 |
| G3 | $\phi_X^G$ | Minimal fraction of supporting fragment ends in contig | 0.5 |

## 3 Pipeline steps

### 3.1 Clustering seed anchors

All contigs that are at least 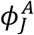 long are clustered using hierarchical clustering, using the shared neighbors metric *D* as a measure of similarity, and using mean linkage for merging clusters. The resulting hierarchical tree is traversed from the root, stopping at the first node *v* that satisfies 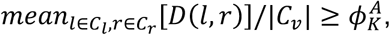 where *C_v_* are the contigs in the sub-tree under the node *v*, and *C_l_*, *C_r_* are the contigs in the sub-trees under the descendants of *v*. The seed anchors are defined as 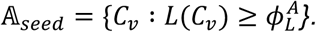

### 3.2 Inference of seed spurious model

A seed model *P* is inferred over all inter-anchor fragment end pairs in 𝔸*_seed_*. The empirical matrix *F_len_* is inferred using maximum likelihood from the data, using all pairs of fragment ends that belong to different seed anchors.

### 3.3 Removing multi-anchor contigs

We denote by *A*(*c*) the anchor of contig c. The greedy algorithm **TrimAnchors** reduces the seed anchors 𝔸_seed_, by removing multi-anchored contigs and updating *P*, until convergence.

#### TrimAnchors(𝔸)

1. Find a contig *c_m_* ∈∪ 𝔸 and an anchor *A_m_* ∈ 𝔸 that satisfy: 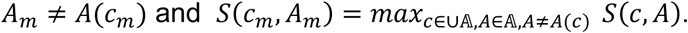
2. If 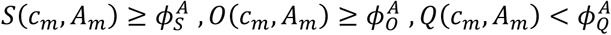 then remove *c_m_* from *A*(*c_m_*).
3. Remove any contig *c* from *A*(*c*) if 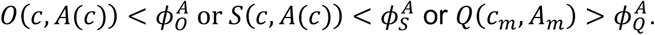
4. Remove any anchor 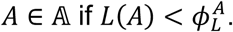
5. Repeat steps 1-4 until no more contigs or anchors are removed.
6. Return 𝔸.

### 3.4 Abundance trimming

Each resulting anchor *A* ∈ 𝔸 is further refined to satisfy the abundance condition (A5) as follows. Any contig *c* ∈ *A* for which 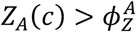 is discarded from *A*. Then the anchor itself is discarded as a whole if it becomes too short 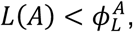 or if the weighted mean standard deviation of the anchor supersedes the threshold 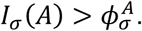

### 3.5 Final model and genome unions

The final model *P* is inferred over the resulting anchor collection 𝔸. For every anchor *A* ∈ 𝔸, the union *G*(*A*) is then computed, to include all contigs c ∈ M that satisfy all of the genome union conditions (G1-G3).

---

**Figure S1.**
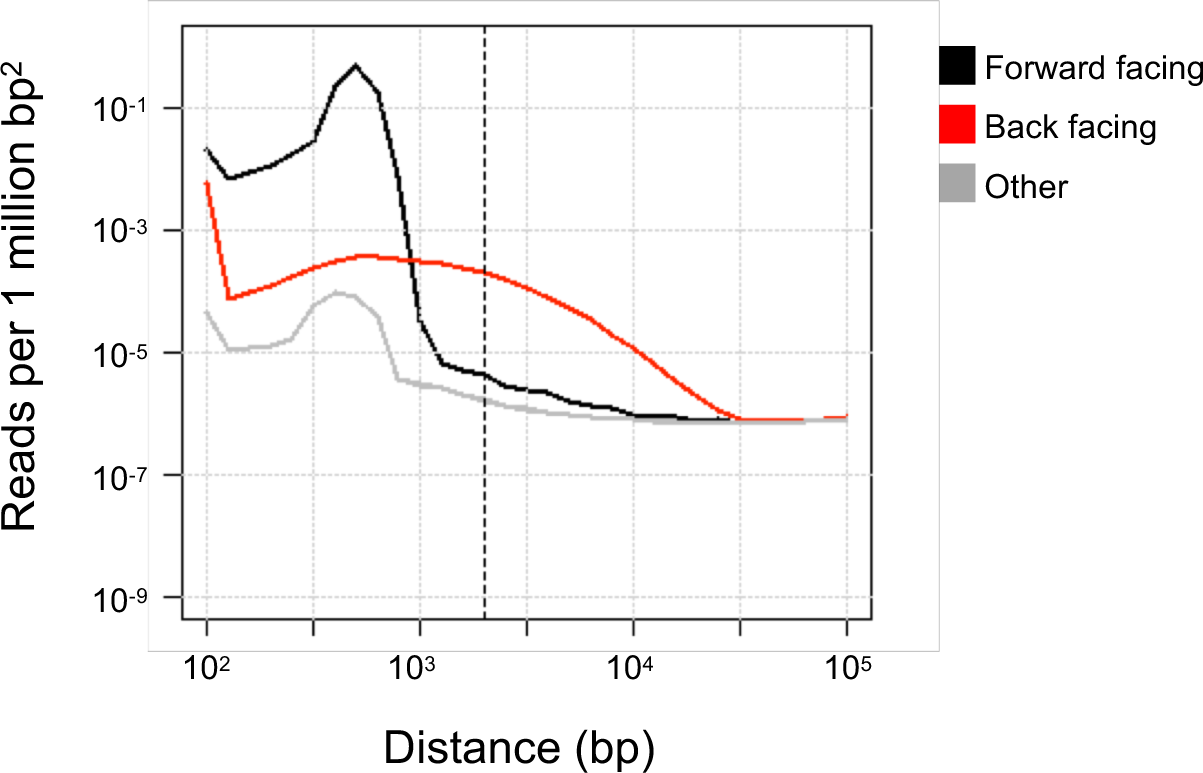
Intra-contig read density as a function distance between mapped read sides, colored according the relative strand orientation of the two read sides.

**Figure S2:**
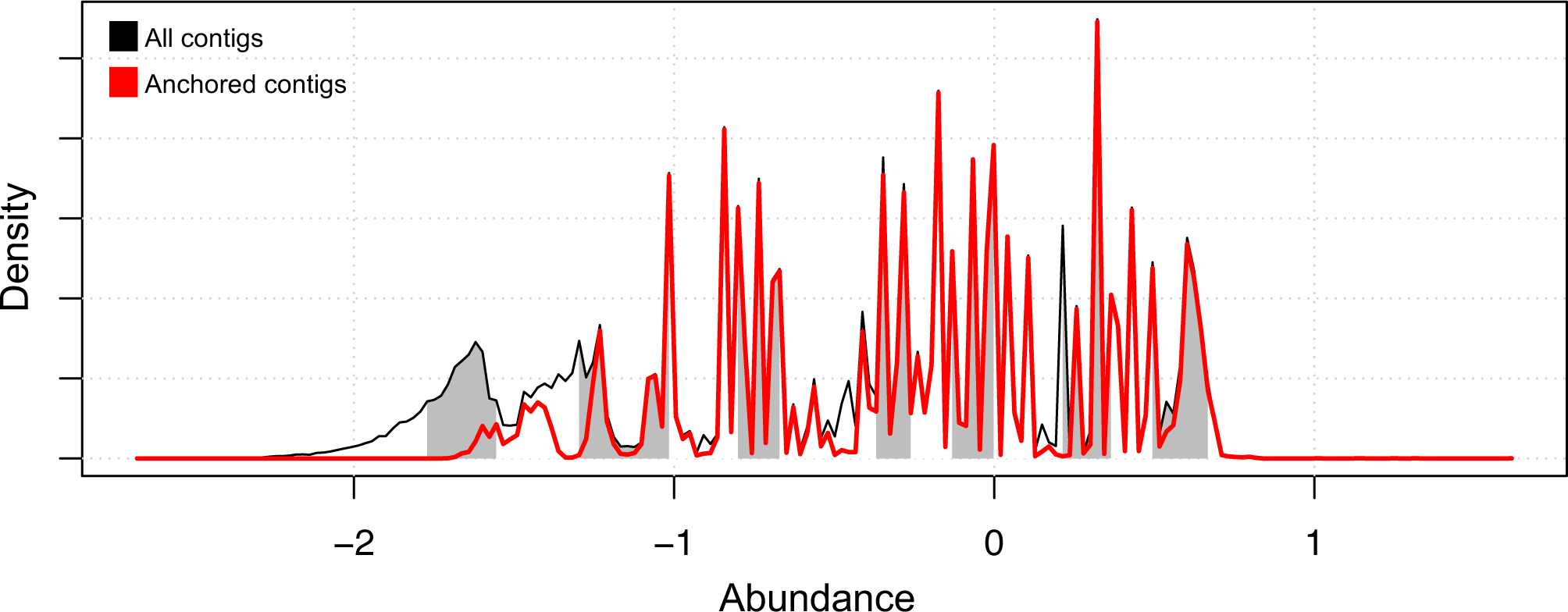
Simulated community. The genomes of 55 common gut microbes (GOLD database) were downloaded and simulated 120M shotgun reads and 100M Hi-C reads were generated, with relative representation ranging from 1 to 1000. HPIPE identified 32 genomes. Shown is the density plot of the relative abundance of the entire metagenomic assembly (contigs >1k), as in Figure 1d. The abundance is the enrichment of the read coverage over a uniform distribution of reads. White/gray stripes denote chunks of 10Mb. The fraction of the assembly that was included in any recovered genome (‘anchored contigs’) is depicted with a red line.

**Figure S3:**
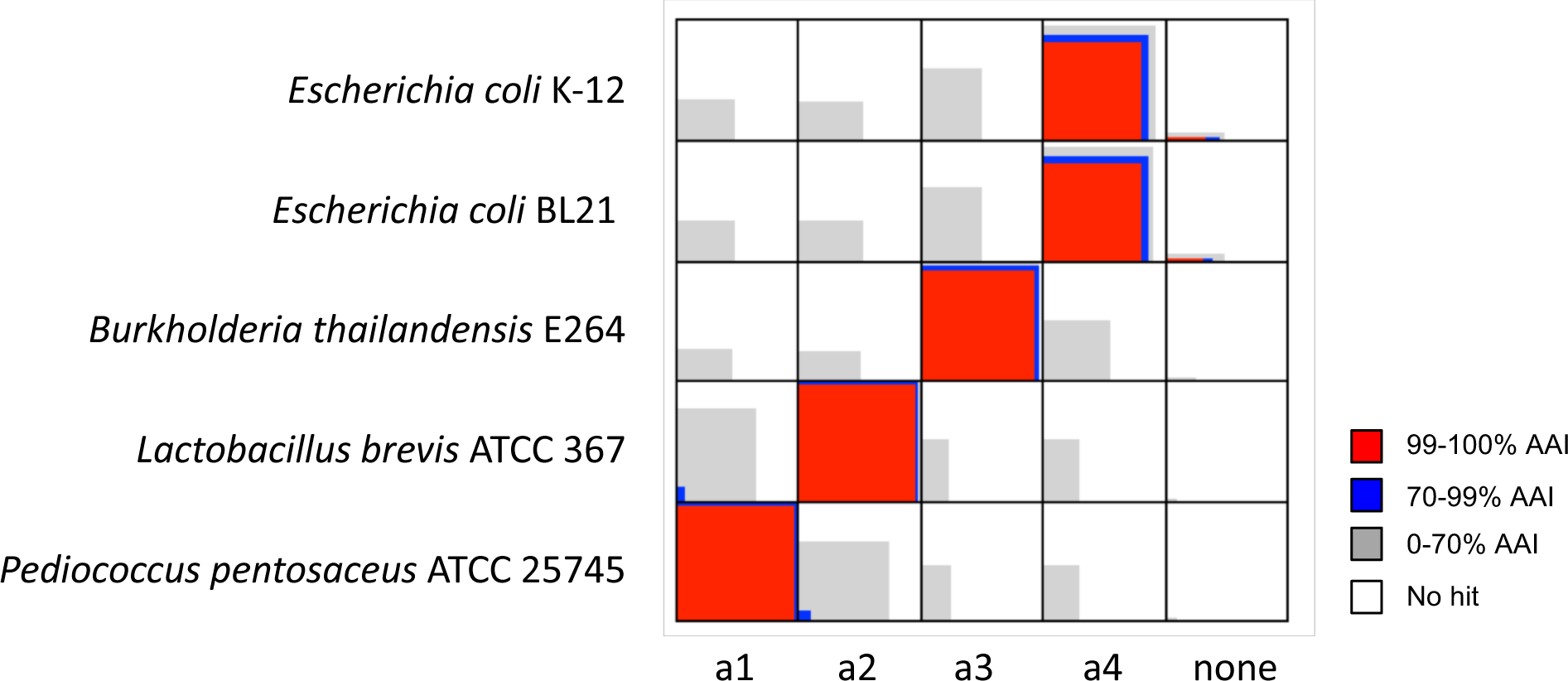
Synthetic community. The community was composed of *Pediococcus pentosaceus* (ATCC 25745), *Lactobacillus brevis* (ATCC 367), *Burkholderia thailandensis* (E264) and two strains of *Escherichia coli* (BL21 and K-12), as described in Beitel et al. 2014. The pipeline recovered 4 anchor/union pairs. Shown is a pairwise gene alignment between the 4 inferred genome unions and the 5 reference genome.

**Figure S4:**
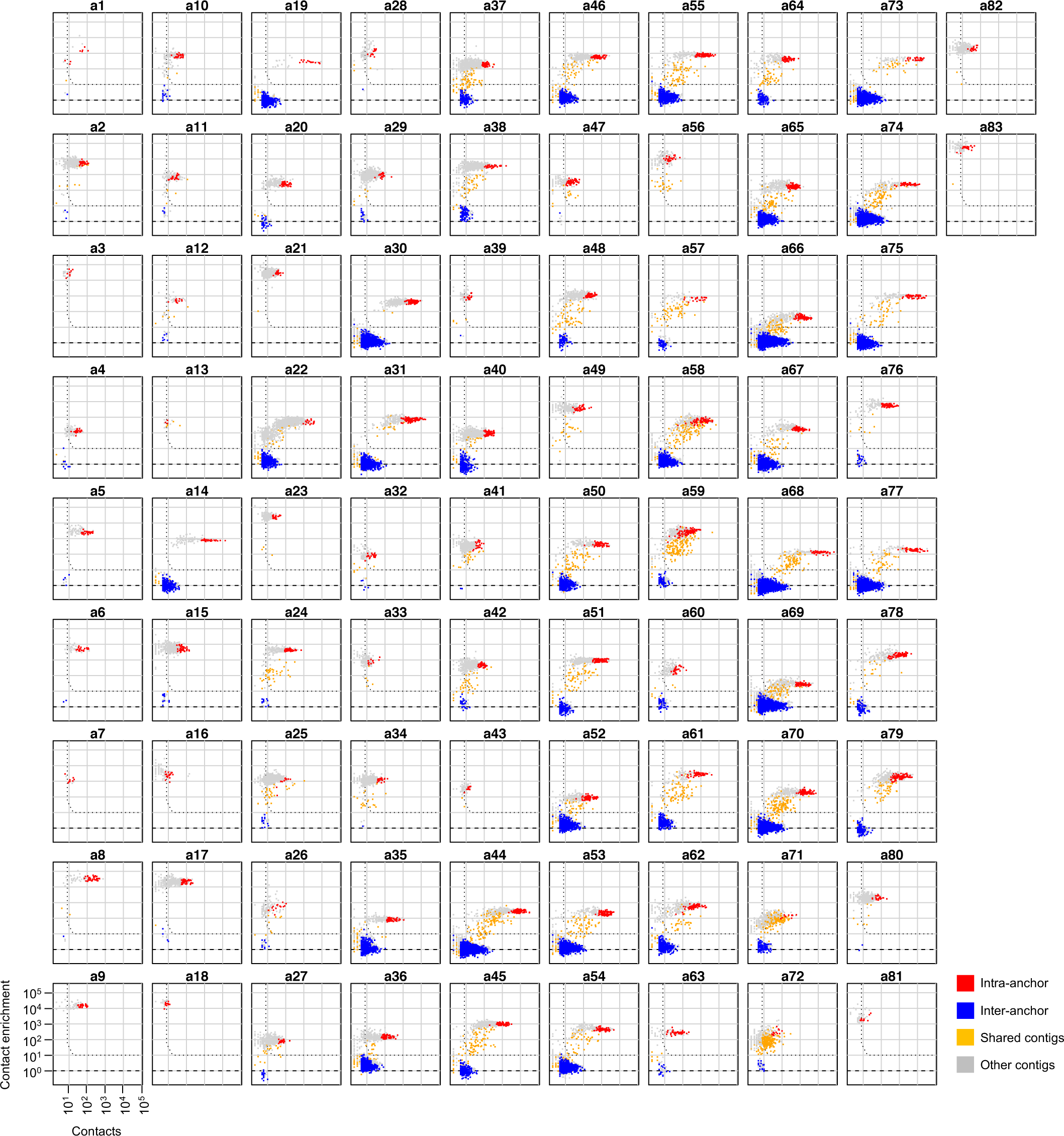
Contig-anchor contact enrichments over all anchors. On the x-axis is the observed number of contacts between the contig and the anchor, and on the y-axis is the enrichment scores over the background model. Anchor contigs are colored red, contigs belonging to other anchors are colored blue, and all other contigs are colored gray. Anchors are extended into genomes by including contigs with >=10-fold contact enrichment (dashed horizontal line), >=8 contacts (dashed vertical line), and a false positive probability of 10^−6^ assuming a binomial distribution (transition between vertical and horizontal line).

**Figure S5:**
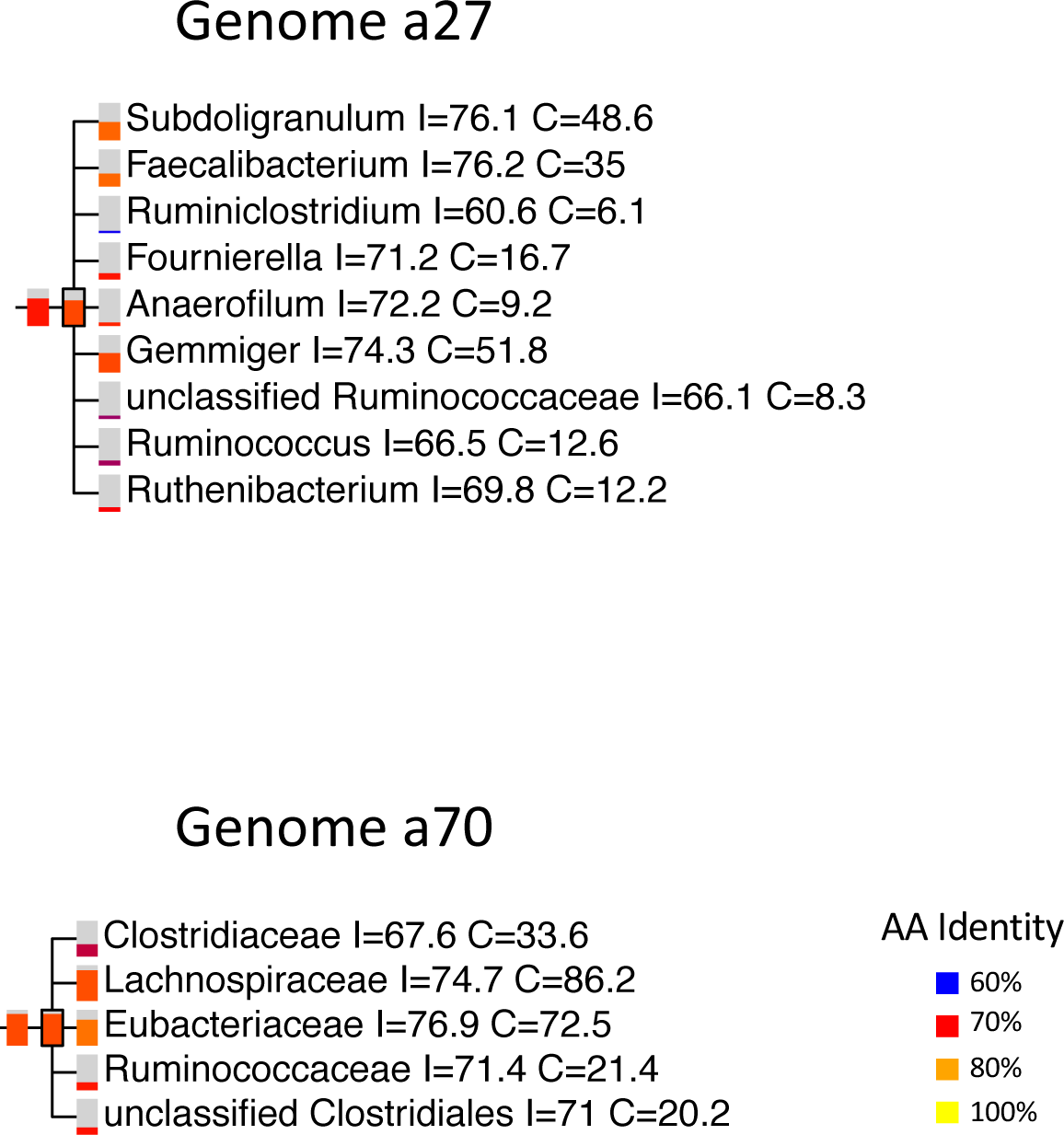
Examples of 2 putative novel genomes. On top, 68% of the genes of genome a27 align to the *Ruminococcaceae* family (mean identity 74.3%), suggesting it is a novel species in that family. On the bottom, 88% of the genes of genome a70 align to the *Clostrdiales* order (mean identity 74.5%), indicating it is a novel genomes under *Lachnospiraceae* or *Eubacteriaceae*. Each taxa is colored according to the mean amino acid identity, and the fraction of colored rectangle represents the percentage of the aligned genes.

**Figure S6:**
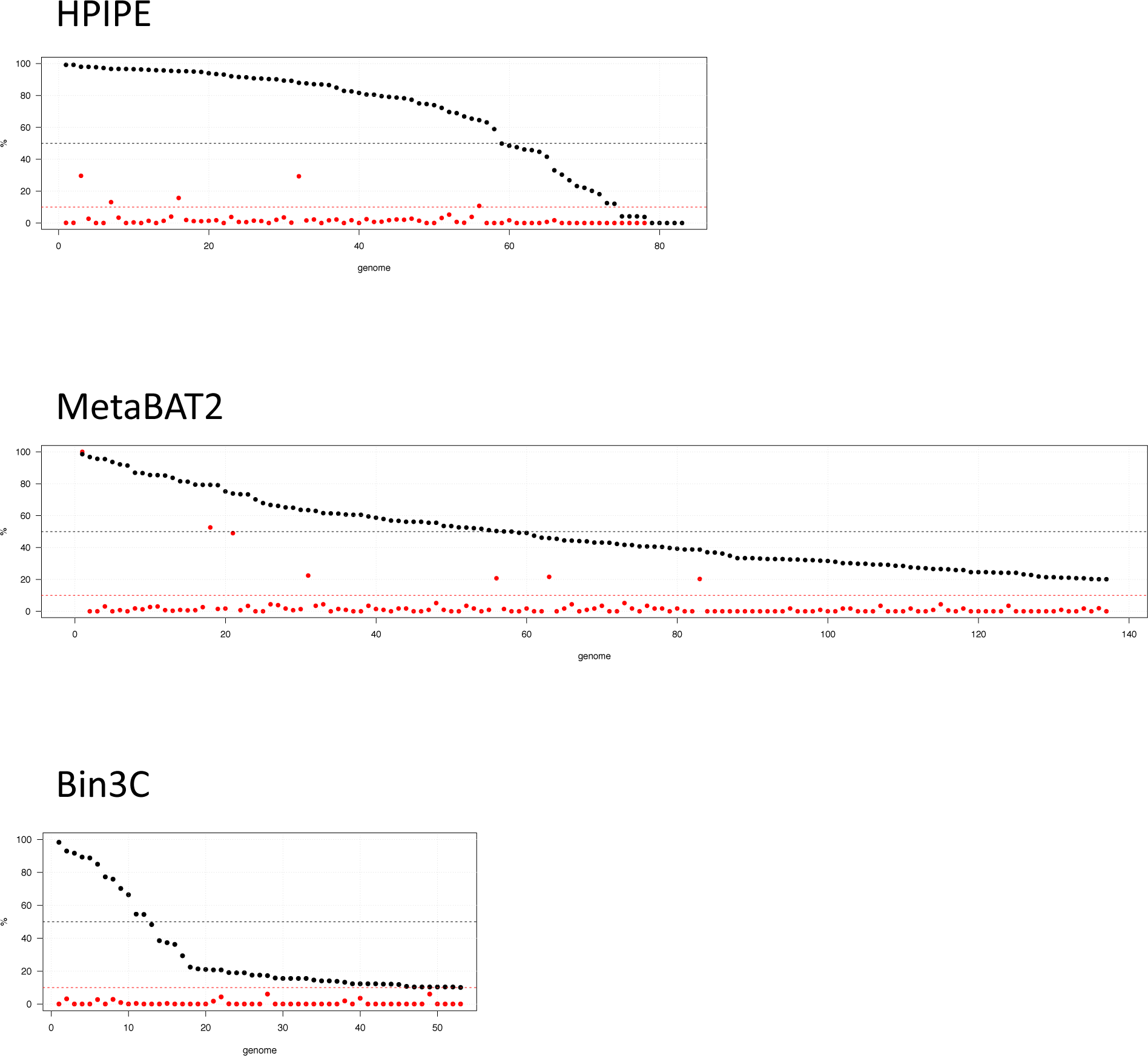
Comparison to alternative metagenomic binning methods. Single-copy gene estimates of genome completeness percentage (in black) and contamination percentage (in red), and sorted according to completeness. Minimal completeness (50%) and maximal contamination (10%) thresholds depicted with dashed horizontal lines. Our results (HPIPE, as in Figure 2c), compared to metaBAT2 (tool based on abundance and tetranucleotide frequency), and bin3C (tool based clustering of Hi-C data).

**Figure S7:**
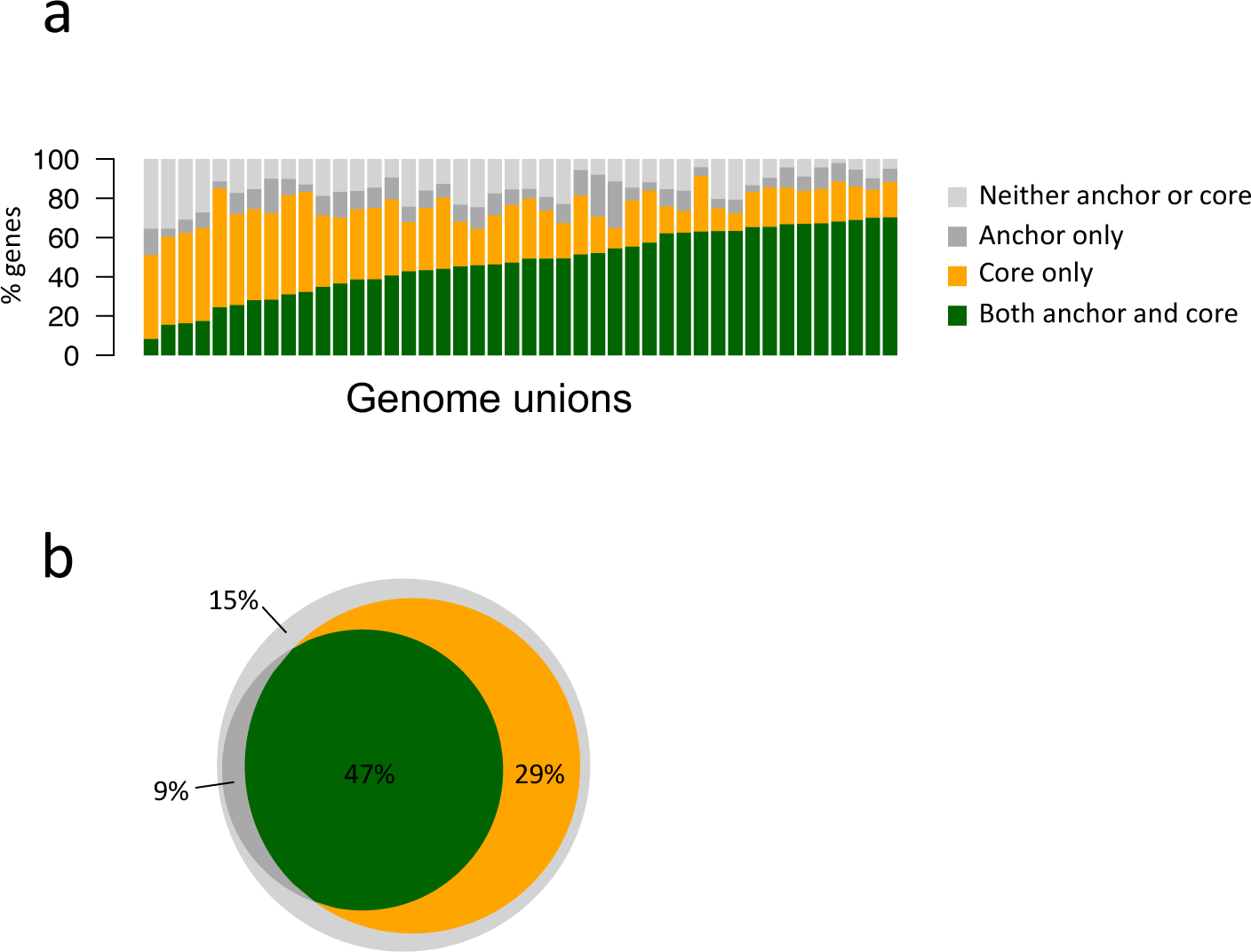
Comparison of anchors and cores. **(a)** Shown for all 44 genome unions, is the breakdown of genes into ‘core-only, ‘anchor-only’, ‘both’ or ‘neither’, sorted according to the ‘both’ fraction. **(b)** The fraction of the 4 gene classifications, colored as in (a), averaged over all 44 genomes. Core-only genes (29%) are present due to the stringent selection of anchors, which considers only long contigs (>10k).

**Figure S8:**
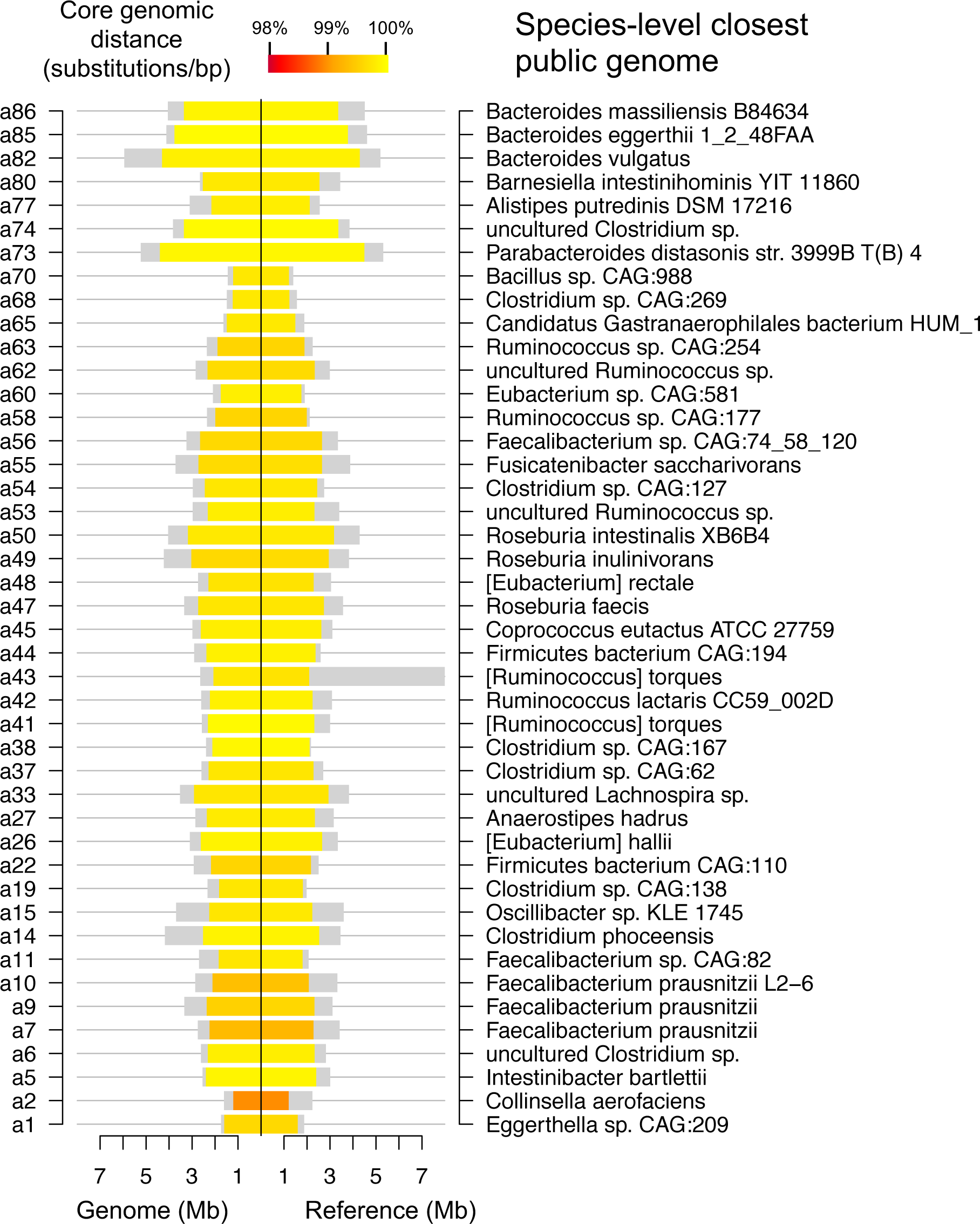
Species-level reference genomes for Subject B. See Figure 3 for legend.

**Figure S9:**
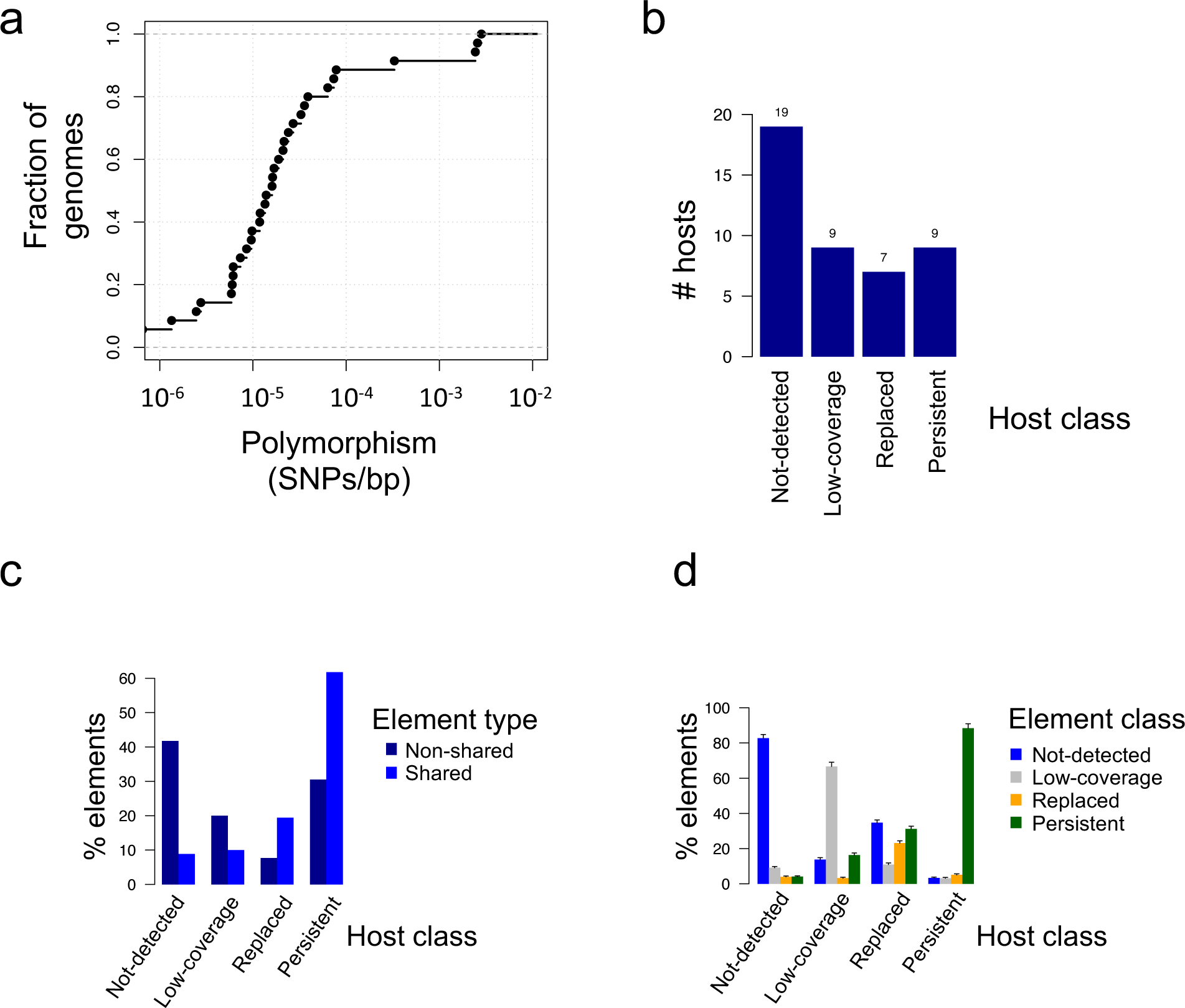
Polymorphism and 10-year divergence patterns for Subject B. **(a)** Polymorphism levels, estimated using the density of intermediate alleles (SNPs with a frequency in the range 20%-80%), shown for 35 genomes of Subject B that had at least 10x coverage. **(b)** Host classification for the 44 genomes of Subject B. **(c-d)** Same analysis as 5c-d, done Subject B.

**Supplementary Table 1.** Gut genomes used for simulated data. 55 bacteria associated with the gut microbiome were downloaded from the GOLD database. Shown for each genome are the species and strain NCBI taxonomic identifier, the taxonomic name and the genome accession identifier.

| index | name | strain.id | species.id | accession |
| --- | --- | --- | --- | --- |
| 1 | Escherichia coli str. K-12 substr. MG1655 | 511145 | 562 | GCA_000005845.2 |
| 2 | Bacteroides vulgatus ATCC 8482 | 435590 | 821 | GCA_000012825.1 |
| 3 | Lactobacillus gasseri ATCC 33323 = JCM 1131 | 324831 | 1596 | GCA_000014425.1 |
| 4 | Lactobacillus ruminis ATCC 27782 | 1069534 | 1623 | GCA_000224985.1 |
| 5 | Bifidobacterium bifidum S17 | 883062 | 1681 | GCA_000164965.1 |
| 6 | Bifidobacterium breve ACS-071-V-Sch8b | 866777 | 1685 | GCA_000213865.1 |
| 7 | Ruminococcus bromii | 40518 | 40518 | GCA_002834165.1 |
| 8 | Roseburia intestinalis L1-82 | 536231 | 166486 | GCA_000156535.1 |
| 9 | Lactobacillus saerimneri 30a | 1227363 | 228229 | GCA_000317165.1 |
| 10 | Clostridium sp. SS2/1 | 411484 | 411484 | GCA_000154545.1 |
| 11 | Clostridium sp. M62/1 | 411486 | 411486 | GCA_000159055.1 |
| 12 | Clostridium sp. L2-50 | 411489 | 411489 | GCA_000154245.1 |
| 13 | Bacteroides sp. 1_1_30 | 457387 | 457387 | GCA_000218365.1 |
| 14 | Parabacteroides sp. 2_1_7 | 457388 | 457388 | GCA_000157035.2 |
| 15 | Bacteroides sp. 3_1_13 | 457389 | 457389 | GCA_001185845.1 |
| 16 | Bacteroides sp. 3_1_23 | 457390 | 457390 | GCA_000162555.1 |
| 17 | Bacteroides sp. 3_2_5 | 457392 | 457392 | GCA_000159855.2 |
| 18 | Bacteroides sp. 4_1_36 | 457393 | 457393 | GCA_000185585.1 |
| 19 | Bacteroides sp. 4_3_47FAA | 457394 | 457394 | GCA_000158515.2 |
| 20 | Bacteroides sp. 9_1_42FAA | 457395 | 457395 | GCA_000157075.2 |
| 21 | Clostridium sp. 1_1_41A1FAA | 457397 | 457397 | GCA_001078415.1 |
| 22 | Synergistes sp. 3_1_syn1 | 457415 | 457415 | GCA_000238615.1 |
| 23 | Bacteroides sp. 1_1_14 | 469585 | 469585 | GCA_000162515.1 |
| 24 | Bacteroides sp. 2_2_4 | 469590 | 469590 | GCA_000157055.1 |
| 25 | Parabacteroides sp. 20_3 | 469591 | 469591 | GCA_000162535.1 |
| 26 | Bacteroides sp. 3_1_40A | 469593 | 469593 | GCA_000186105.1 |
| 27 | Bacteroides sp. D1 | 556258 | 556258 | GCA_000157095.2 |
| 28 | Bacteroides sp. D2 | 556259 | 556259 | GCA_000159075.2 |
| 29 | Parabacteroides sp. D13 | 563193 | 563193 | GCA_000162275.1 |
| 30 | Bacteroides sp. D20 | 585543 | 585543 | GCA_000162215.1 |
| 31 | Lachnospiraceae bacterium 6_1_63FAA | 658083 | 658083 | GCA_000209425.1 |
| 32 | Parabacteroides sp. D26 | 658662 | 658662 | GCA_001078555.1 |
| 33 | Porphyromonas sp. 31_2 | 658663 | 658663 | GCA_000712235.1 |
| 34 | Campylobacter sp. 10_1_50 | 665939 | 665939 | GCA_000238755.1 |
| 35 | Clostridium sp. 7_3_54FAA | 665940 | 665940 | GCA_000233515.1 |
| 36 | Tannerella sp. 6_1_58FAA_CT1 | 665949 | 665949 | GCA_000238695.1 |
| 37 | Subdoligranulum sp. 4_3_54A2FAA | 665956 | 665956 | GCA_000238635.1 |
| 38 | Bacillus sp. 7_6_55CFAA_CT2 | 665957 | 665957 | GCA_000238655.1 |
| 39 | Bacillus sp. BT1B_CT2 | 665958 | 665958 | GCA_000186125.1 |
| 40 | Bilophila sp. 4_1_30 | 693988 | 693988 | GCA_000224655.1 |
| 41 | Lactobacillus rogosae | 706562 | 706562 | GCA_900112995.1 |
| 42 | Rahnella sp. Y9602 | 741091 | 741091 | GCA_000187705.1 |
| 43 | Collinsella sp. 4_8_47FAA | 742722 | 742722 | GCA_000763055.1 |
| 44 | Coproccoccus sp. ART55/1 | 751585 | 751585 | GCA_000210595.1 |
| 45 | Clostridium sp. HGF2 | 908340 | 908340 | GCA_000183585.2 |
| 46 | Paenibacillus sp. HGF5 | 908341 | 908341 | GCA_000204455.2 |
| 47 | Alistipes sp. HGB5 | 908612 | 908612 | GCA_000183485.2 |
| 48 | Paenibacillus sp. HGF7 | 944559 | 944559 | GCA_000214295.2 |
| 49 | Brevibacterium senegalense | 1033736 | 1033736 | GCA_000285835.2 |
| 50 | Kurthia massiliensis | 1033739 | 1033739 | GCA_000285555.1 |
| 51 | Dielma fastidiosa | 1034346 | 1034346 | GCA_000313565.2 |
| 52 | Alistipes obesi | 1118061 | 1118061 | GCA_000311925.1 |
| 53 | Verrucomicrobia bacterium SCGC AB-629-E09 | 1131271 | 1131271 | GCA_000371985.1 |
| 54 | Ruminococcus bicirculans | 1160721 | 1160721 | GCA_000723465.1 |
| 55 | Megasphaera massiliensis | 1232428 | 1232428 | GCA_000455225.1 |

**Supplementary Table 2.**
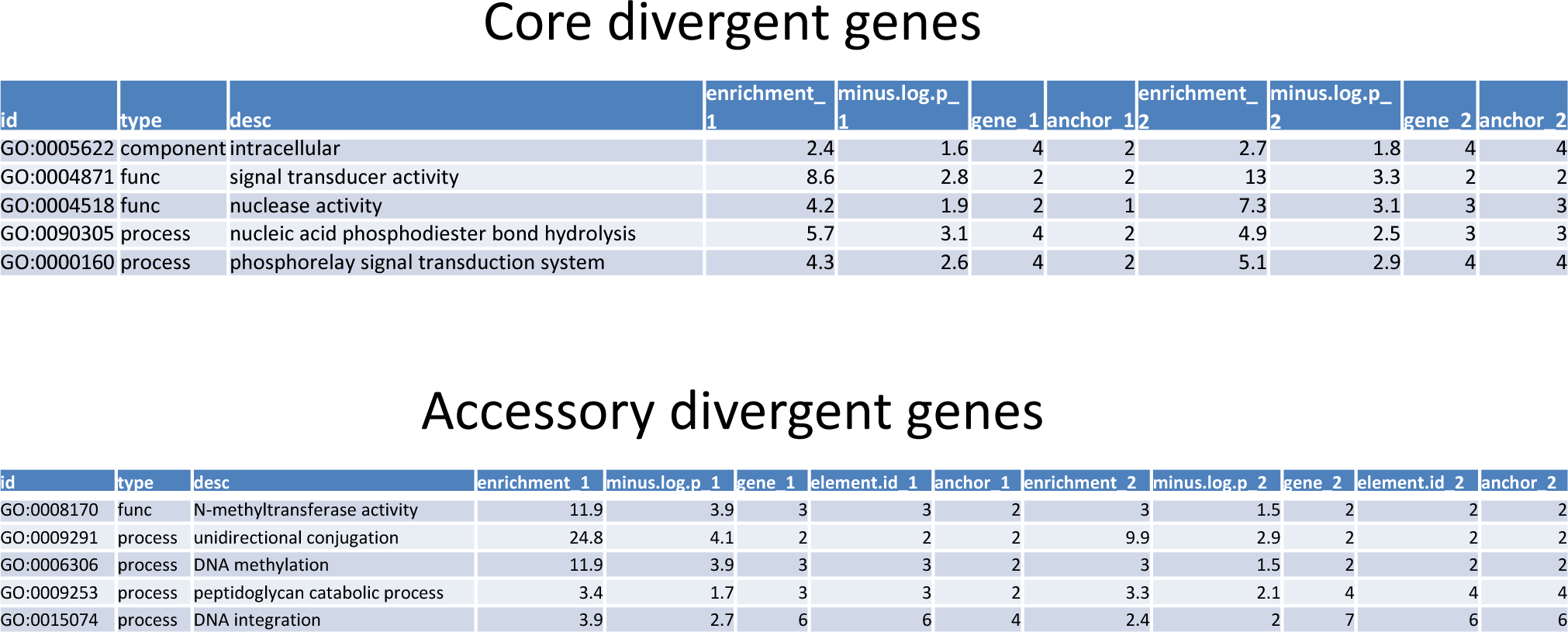
Gene Ontology for evolved genes. GO annotations shown for 152 genes that contained non-synonymous substitutions (core divergent genes, top), and for the 1253 accessory genes that resided on non-persistent accessory elements that were associated with persistent hosts (accessory divergent genes, bottom). Shown for each GO category is the enrichment (fold-change of gene count above background), the chi-square p-value (hypergeometric test), the number of genes, unique elements and unique associated hosts, separately for both subjects (Subject A and B have a suffix of ‘_1’ and ‘_2’ respectively). The background used for all tests was the entire set of predicted genes.

Core divergent genes
| id | type | desc | enrichment_1 | minus.log.p_1 | gene_1 | anchor_1 | enrichment_2 | minus.log.p_2 | gene_2 | anchor_2 |
| --- | --- | --- | --- | --- | --- | --- | --- | --- | --- | --- |
| GO:0005622 | component | intracellular | 2.4 | 1.6 | 4 | 2 | 2.7 | 1.8 | 4 | 4 |
| GO:0004871 | func | signal transducer activity | 8.6 | 2.8 | 2 | 2 | 13 | 3.3 | 2 | 2 |
| GO:0004518 | func | nuclease activity | 4.2 | 1.9 | 2 | 1 | 7.3 | 3.1 | 3 | 3 |
| GO:0090305 | process | nucleic acid phosphodiester bond hydrolysis | 5.7 | 3.1 | 4 | 2 | 4.9 | 2.5 | 3 | 3 |
| GO:0000160 | process | phosphorelay signal transduction system | 4.3 | 2.6 | 4 | 2 | 5.1 | 2.9 | 4 | 4 |

**Supplementary Table 2. Gene Ontology for evolved genes.**
| id | type | desc | enrichment_1 | minus.log.p_1 | gene_1 | element.id_1 | anchor_1 | enrichment_2 | minus.log.p_2 | gene_2 | element.id_2 | anchor_2 |
| --- | --- | --- | --- | --- | --- | --- | --- | --- | --- | --- | --- | --- |
| GO:0008170 | func | N-methyltransferase activity | 11.9 | 3.9 | 3 | 3 | 2 | 3 | 1.5 | 2 | 2 | 2 |
| GO:0009291 | process | unidirectional conjugation | 24.8 | 4.1 | 2 | 2 | 2 | 9.9 | 2.9 | 2 | 2 | 2 |
| GO:0006306 | process | DNA methylation | 11.9 | 3.9 | 3 | 3 | 2 | 3 | 1.5 | 2 | 2 | 2 |
| GO:0009253 | process | peptidoglycan catabolic process | 3.4 | 1.7 | 3 | 3 | 2 | 3.3 | 2.1 | 4 | 4 | 4 |
| GO:0015074 | process | DNA integration | 3.9 | 2.7 | 6 | 6 | 4 | 2.4 | 2 | 7 | 6 | 6 |

**Supplementary Table 3.**
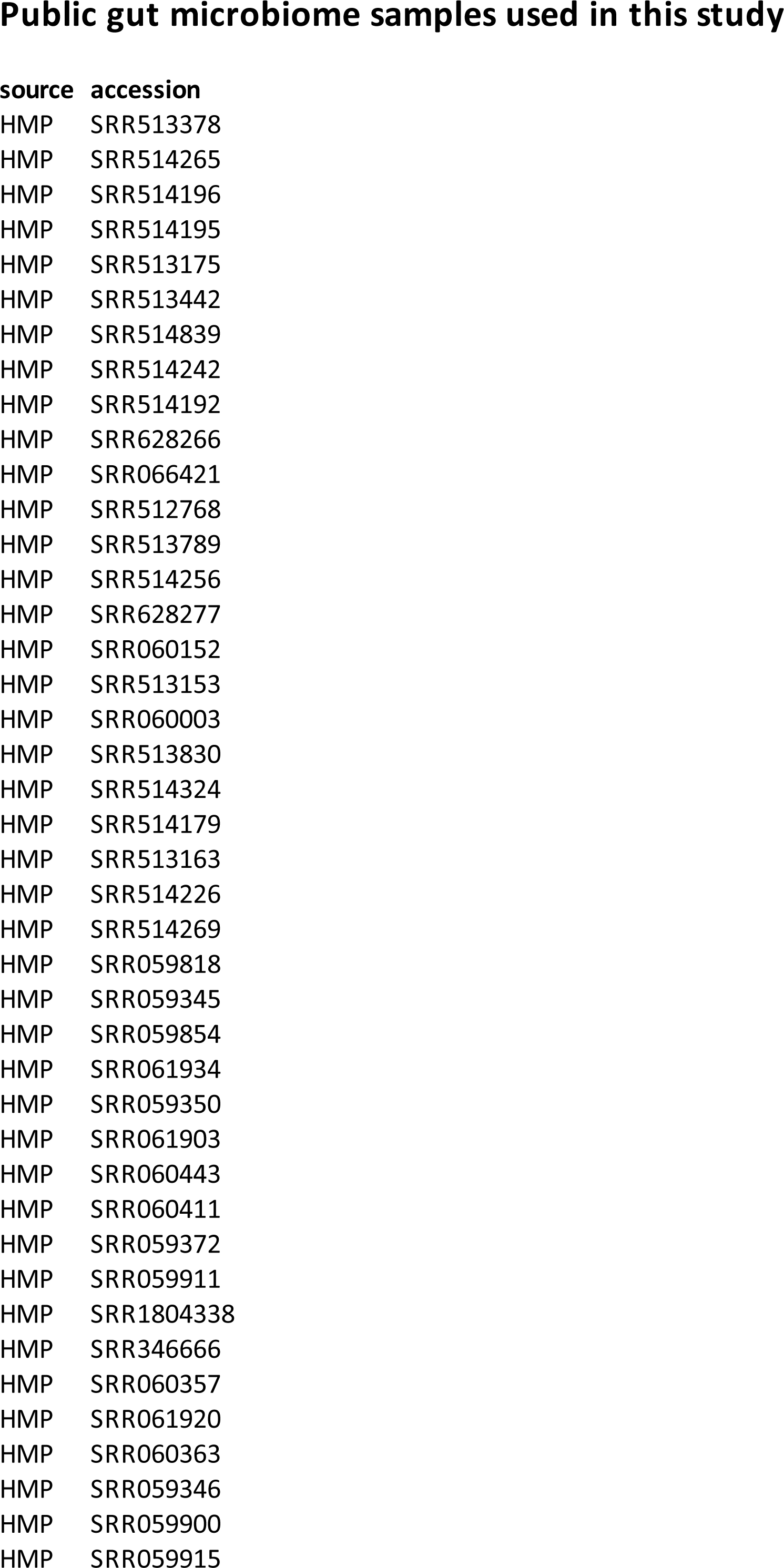
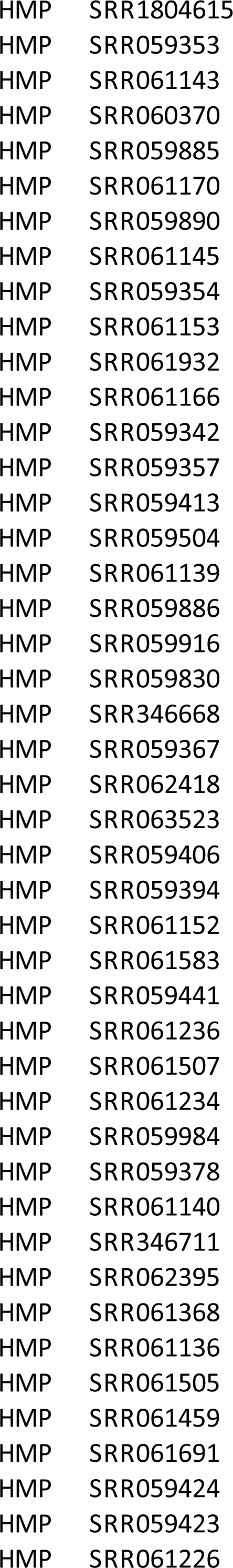
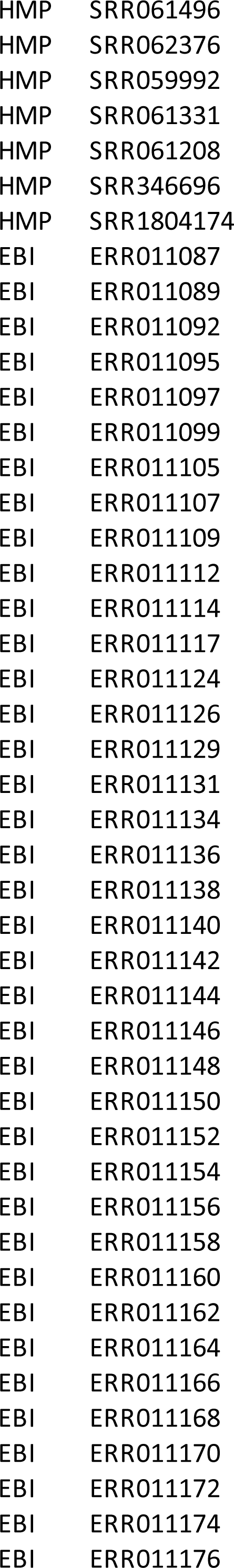
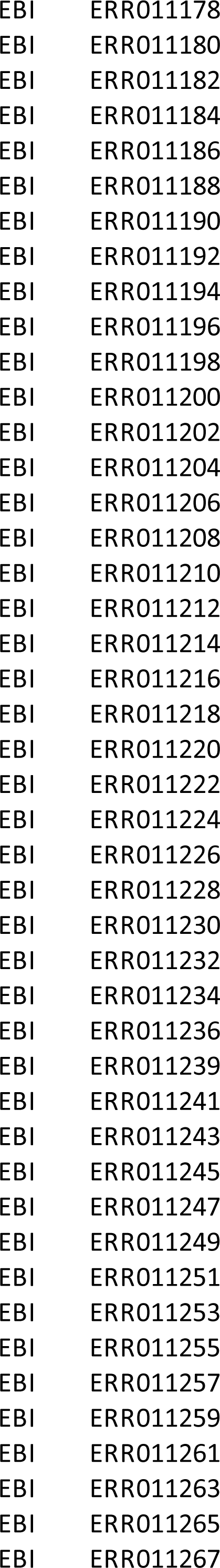
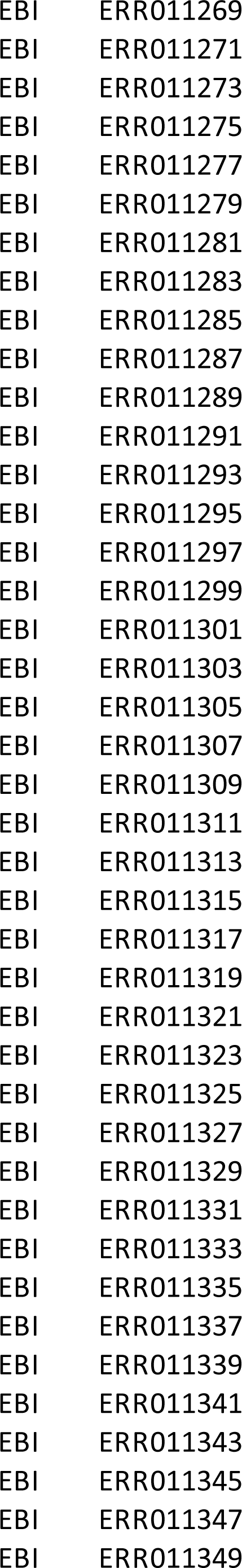
Public gut microbiome samples used in this study. Shown for all 218 datasets are the source (HMP or EBI) and the accession identifier.

**Supplementary Table 4.**
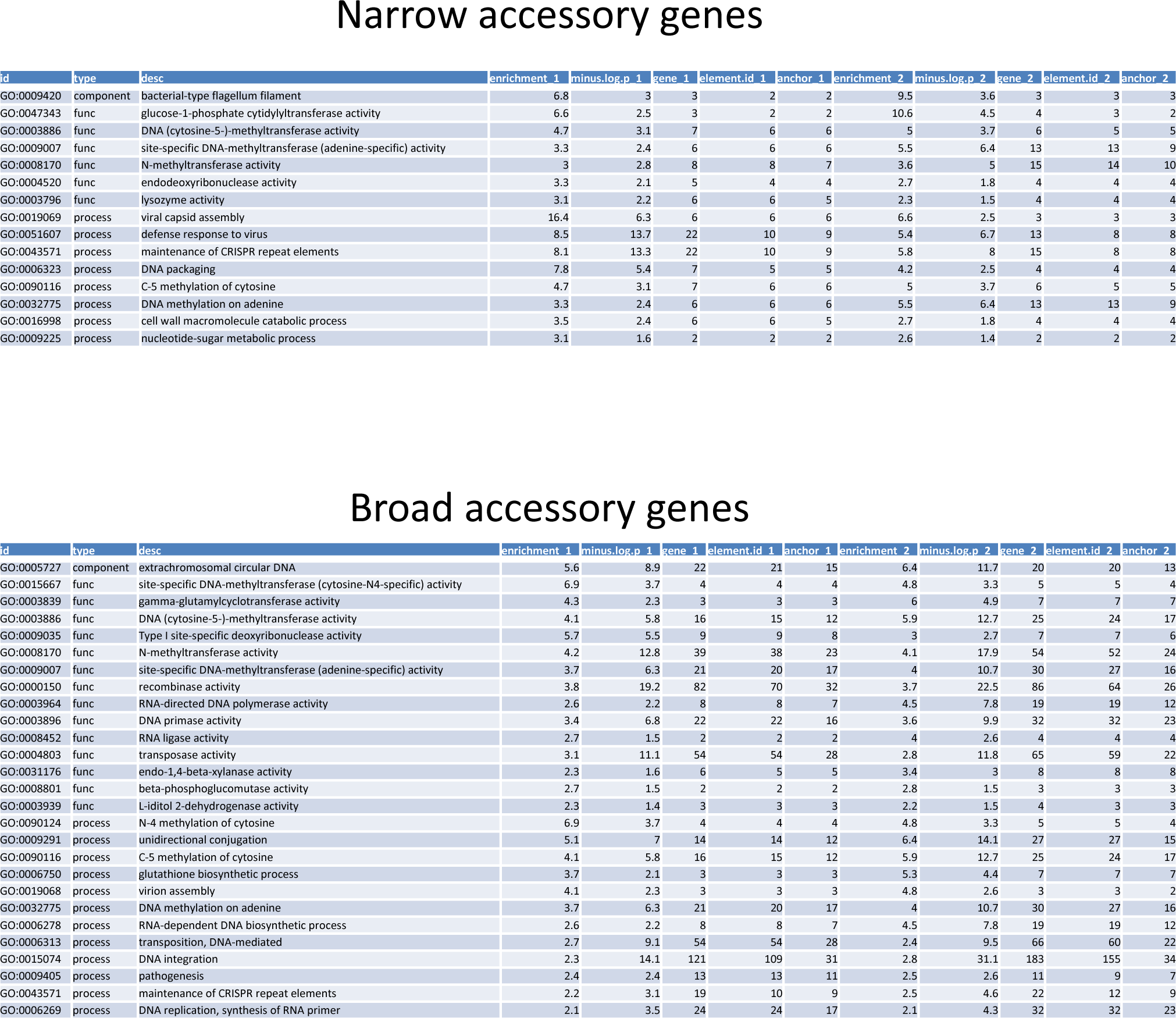
Gene Ontology for narrow-range and broad-range elements. GO annotations shown for genes residing on narrow-range elements (narrow accessory genes, top), and for genes residing on broad-range elements (broad accessory genes, bottom). Shown for each GO category is the enrichment (fold-change of gene count above background), the chi-square p-value (hypergeometric test), the number of genes, unique elements and unique associated hosts, separately for both subjects (Subject A and B have a suffix of ‘_1’ and ‘_2’ respectively). The background used for all tests was the entire set of predicted genes.

